# A potential patient stratification biomarker for Parkinso’s disease based on LRRK2 kinase-mediated centrosomal alterations in peripheral blood-derived cells

**DOI:** 10.1101/2023.04.11.536367

**Authors:** Yahaira Naaldijk, Belén Fernández, Rachel Fasiczka, Elena Fdez, Coline Leghay, Ioana Croitoru, John B. Kwok, Yanisse Boulesnane, Amelie Vizeneux, Eugenie Mutez, Camille Calvez, Alain Destée, Jean-Marc Taymans, Ana Vinagre Aragon, Alberto Bergareche Yarza, Shalini Padmanabhan, Mario Delgado, Roy N. Alcalay, Zac Chatterton, Nicolas Dzamko, Glenda Halliday, Javier Ruiz-Martínez, Marie-Christine Chartier-Harlin, Sabine Hilfiker

## Abstract

Parkinso’s disease (PD) is a common neurodegenerative movement disorder and leucine-rich repeat kinase 2 (LRRK2) is a promising therapeutic target for disease intervention. However, the ability to stratify patients who will benefit from such treatment modalities based on shared etiology is critical for the success of disease-modifying therapies. Ciliary and centrosomal alterations are commonly associated with pathogenic LRRK2 kinase activity and can be detected in many cell types. We previously found centrosomal deficits in immortalized lymphocytes from *G2019S-LRRK2* PD patients. Here, to investigate whether such deficits may serve as a potential blood biomarker for PD which is susceptible to LRKK2 inhibitor treatment, we characterized patient-derived cells from distinct PD cohorts. We report centrosomal alterations in peripheral cells from a subset of early-stage idiopathic PD patients which is mitigated by LRRK2 kinase inhibition, supporting a role for aberrant LRRK2 activity in idiopathic PD. Centrosomal defects are detected in *R1441G-LRRK2* and *G2019S-LRRK2* PD patients and in non-manifesting *LRRK2* mutation carriers, indicating that they acumulate prior to a clinical PD diagnosis. They are present in immortalized cells as well as in primary lymphocytes from peripheral blood. These findings indicate that analysis of centrosomal defects as a blood-based patient stratification biomarker may help nominate PD patients who will benefit from LRRK2-related therapeutics.

**One-sentence summary:** Peripheral blood-derived cells can be employed to stratify Parkinso’s disease patients most likely to respond to LRRK2-related therapeutics.

## Introduction

Parkinso’s disease (PD) is characterized by the progressive loss of dopaminergic neurons in the substantia nigra which results in motor symptoms such as tremor, rigidity, bradykinesia and postural instability. At the time of clinical diagnosis, a large percentage of these neurons have already degenerated (*1*). Whilst current therapies can temporarily improve motor symptoms, there are no treatments which slow or halt the disease. PD patients display differences in clinical symptoms and rates of disease progression which may reflect distinct underlying molecular and biological alterations (*2, 3*). Hence, identifying biomarkers for the early diagnosis of at least some types of PD and for the assessment of therapeutic interventions has become a key challenge in the field.

Genetic variations in *leucine-rich repeat kinase 2 (LRRK2)* are strongly implicated in PD risk. Distinct missense mutations are a frequent cause of autosomal-dominant inherited PD, and common variants in the *LRRK2* gene are associated with a greater risk of developing idiopathic PD (*4–8*). All known familial missense mutations increase the kinase activity of LRRK2 (*9, 10*). Increased kinase activity has also been detected in PD patients with certain other genetic forms of PD and in postmortem brain tissue from at least some idiopathic PD patients (*11–13*). These findings indicate that increased LRRK2 activity may be implicated in a significant portion of PD cases.

LRRK2 is highly expressed in peripheral immune cells as compared to central nervous system (*14*), suggesting that blood-based assays may allow for the identification of PD patients who share pathogenic mechanisms due to elevated LRRK2 activity. Increased LRRK2 kinase activity results in enhanced autophosphorylation and phosphorylation of substrates including Rab GTPases which act as master regulators of intracellular trafficking events (*9, 15–17*). Therefore, expression or phosphorylation levels of LRRK2 and its Rab substrates have the potential to serve as biomarkers for PD due to increased LRRK2 activity (*18*). However, extensive studies in blood-derived cells employing distinct approaches to detect levels/phosphorylation of LRRK2 or Rab substrates have been relatively unsuccessful in differentiating *LRRK2* mutation PD patients or idiopathic PD patients from healthy controls (*18–28*).

Cellular consequences downstream of enhanced LRRK2 kinase activity such as lysosomal dysfunction (*29*), which can lead to lysosomal exocytosis, may comprise alternative LRRK2 biomarkers. Lysobisphosphatidic acid (also called BMP [bis(monoacylglycerol)phosphate]), a phospholipid in late endosomes/lysosomes, is increased in urine in *G2019S-LRRK2* PD cases compared to healthy controls and is currently employed as a biomarker in clinical trials with LRRK2 kinase inhibitors (*30–32*). Similarly, increased LRRK2 autophosphorylation and Rab substrate phosphorylation can be detected in urinary exosomes from *LRRK2* mutation PD patients (*33–37*), but none of these urinary measures reliably stratify idiopathic PD patients who may benefit from LRRK2 inhibitor treatment approaches. Hence, there exists an unmet need for patient stratification biomarkers able to define not only *LRRK2* variant carriers but also subgroups of idiopathic PD patients who share the same LRRK2 kinase-mediated deficits.

Rab10 is a prominent LRRK2 kinase substrate (*9, 16*). Phosphorylation of Rab10 impairs its normal function in membrane trafficking (*38, 39*) but allows it to interact with a new set of effector proteins including RILPL1 (*16*). RILPL1 is localized at the mother centriole and recruits phosphorylated Rab10 to this location (*40, 41*). The mother centriole forms the base upon which cilia are formed, and the centriolar phospho-Rab10/RILPL1 complex blocks cilia formation in a variety of cell types *in vitro* (*16, 40, 42, 43*). Ciliogenesis deficits are also observed in certain neurons and astrocytes in pathogenic *G2019S-LRRK2* or *R1441C-LRRK2* knockin mouse models (*40, 44*), suggesting that they are a direct cellular consequence of pathogenic *LRRK2* mutations and observable in the intact rodent brain.

The LRRK2-mediated centriolar phospho-Rab10/RILPL1 complex plays additional roles in non-ciliated cells. In interphase cells, the mother and daughter centriole associate to form a single centrosome in a process called centriole cohesion. Upon centriole duplication in S phase of the cell cycle, the two centrosomes are held together by a process called centrosome cohesion, and both centriole and centrosome cohesion are mediated by a common set of linker proteins (*45–50*). We have previously shown that mutant LRRK2 causes centrosomal cohesion deficits which are dependent on the presence of RILPL1 and Rab10 in a variety of cell types *in vitro* (*42, 43, 51*). Centrosomal cohesion deficits were further observed in immortalized lymphocytes (LCLs) from a cohort of *G2019S-LRRK2* PD patients as compared to healthy controls, and were reverted by the LRRK2 kinase inhibitor MLi2 in all cases (*52*). MLi2-sensitive cohesion deficits were also present in several early-stage idiopathic PD patients, suggesting the possibility that this cellular readout may help to stratify idiopathic PD patients susceptible to LRRK2-related therapeutics (*52*).

Here, we present evidence that LRRK2 kinase activity-mediated cohesion deficits are common to distinct *LRRK2* mutation carriers, detectable in a subset of idiopathic PD patients and present in peripheral blood-derived cells. Our data substantiate a role for increased LRRK2 kinase activity in at least some idiopathic PD patients and suggest that blood-based PD patient stratification according to cohesion deficits may hold promise in the context of clinical trials with LRRK2 inhibitors.

## Results

### Centrosomal cohesion deficits in a subset of idiopathic PD patient-derived cells are mitigated by LRRK2 kinase inhibition

We first determined whether centrosomal cohesion deficits can be observed in a larger sampling of idiopathic PD patients. For this purpose, we employed control and idiopathic PD patient-derived Epstein-Barr virus (EBV)-transformed lymphoblastoid cell lines (LCLs) from a cohort of PD patients (n=35) and controls (n=3) (**Table 1**). Centrosomes were only scored when positive for two distinct centrosomal markers (γ-tubulin and pericentrin), since the percentage of cells with duplicated centrosomes as determined by such an approach closely matches the percentage of cells in G2 phase as determined by flow cytometry (*52*). We previously determined that the mean distance between duplicated centrosomes in healthy control LCLs was around 1.3 μm, and centrosomes were considered as separated when the distance between them was bigger than this value (*52*). When quantifying the distance between duplicated centrosomes in the different LCL lines, ten out of 35 PD LCLs displayed a centrosomal cohesion deficit (**Fig 1A, B**). This deficit was reverted upon short-term incubation with the LRRK2 kinase inhibitor MLi2 in all cases (**Fig. 1B,C**). It was not associated with changes in the percentage of cells displaying two centrosomes (**Fig. 1D**), and did not correlate with gender or age at diagnosis (not shown).

**Figure 1.**
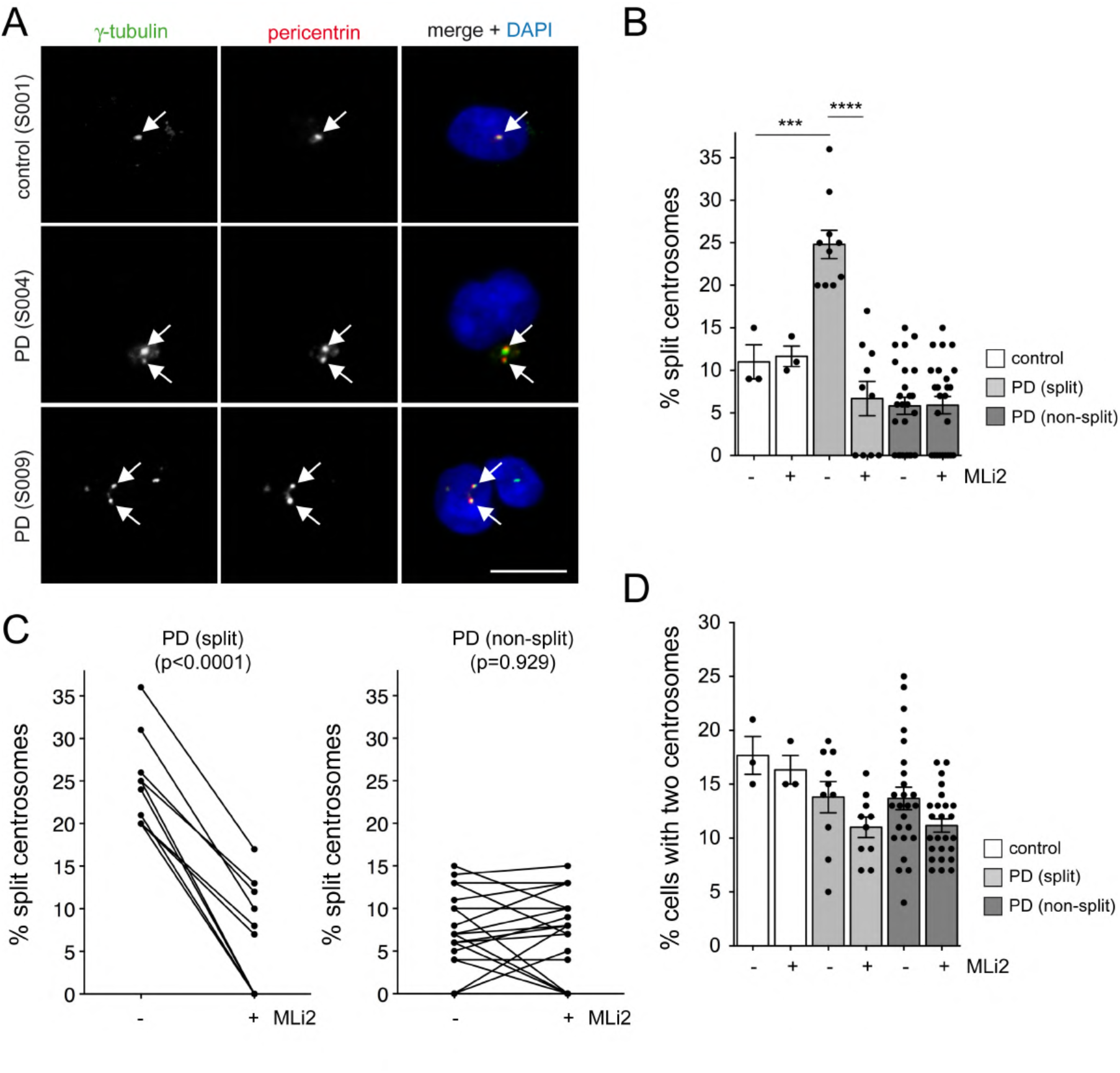
A subset of PD patient LCLs display centrosomal cohesion deficits reverted by short-term treatment with the LRRK2 kinase inhibitor MLi2. (A) Example of one healthy control and two PD LCL lines stained for two centrosomal markers (γ-tubulin and pericentrin) and DAPI. Arrows point to centrosomes co-stained with both markers. Scale bar, 10 μm. (B) The centrosome phenotype was quantified from 100-150 cells per line from 3 control and 35 PD lines, with 10/35 lines (28%) found to display a centrosomal cohesion deficit reverted by MLi2 (50 nM, 2 h). Bars represent mean ± s.e.m.; control versus PD (split) (p = 0.001); PD (split) versus PD (split) + MLi2, (p < 0.001). ***p < 0.005; ****p < 0.001. (C) Paired t-test analysis of centrosomal cohesion deficits from each cell line in the absence or presence of MLi2 as indicated. Note that differences in the values between 0 and 15% are not significant given the small number of cells displaying a duplicated split centrosome phenotype. (D) Quantification of the percent of cells displaying two centrosomes (positive for both pericentrin and γ-tubulin) from a total of 100-150 cells per LCL line.

**Table 1.** Demographic details for Parkinso’s disease and control patient LCLs. Age at diagnosis (mean ± s.d.) and gender of participants is indicated. Apart from a neurological diagnosis (PD versus healthy control), there was limited further clinical information available.

| **Table 1** Demographic details of participants |  |  |
| --- | --- | --- |
|  | Controls | PD |
| Participant number | 3 | 35 |
| Age (y ± s.d.) | 74 ± 15.8 | 65 ± 13.8 |
| Sex (% male) | 33 % | 63 % |

Quantitative immunoblotting of extracts from the different idiopathic PD LCLs showed highly variable levels of total LRRK2, but no differences in the levels of LRRK2 between LCLs with or without a centrosomal cohesion deficit (**Fig. 2A,B**). Similarly, no differences were observed in the levels of pS935-LRRK2, pT73-Rab10 or total Rab10 amongst idiopathic PD LCLs with or without a cohesion deficit (**Fig. 2B**). Correlation analysis was performed to determine possible associations between LRRK2 and pT73-Rab10 levels across all samples. This analysis indicated a significant positive correlation between LRRK2 levels and pT73-Rab10 phosphorylation (**Fig. 2C**). Thus, and similar to what we previously reported for a different cohort of idiopathic PD LCLs (*52*), LRRK2 kinase activity-mediated centrosomal cohesion deficits are detectable in a subset of idiopathic PD samples, even though they do not correlate with increased LRRK2 or pT73-Rab10 levels as assessed by quantitative Western blotting techniques.

**Figure 2.**
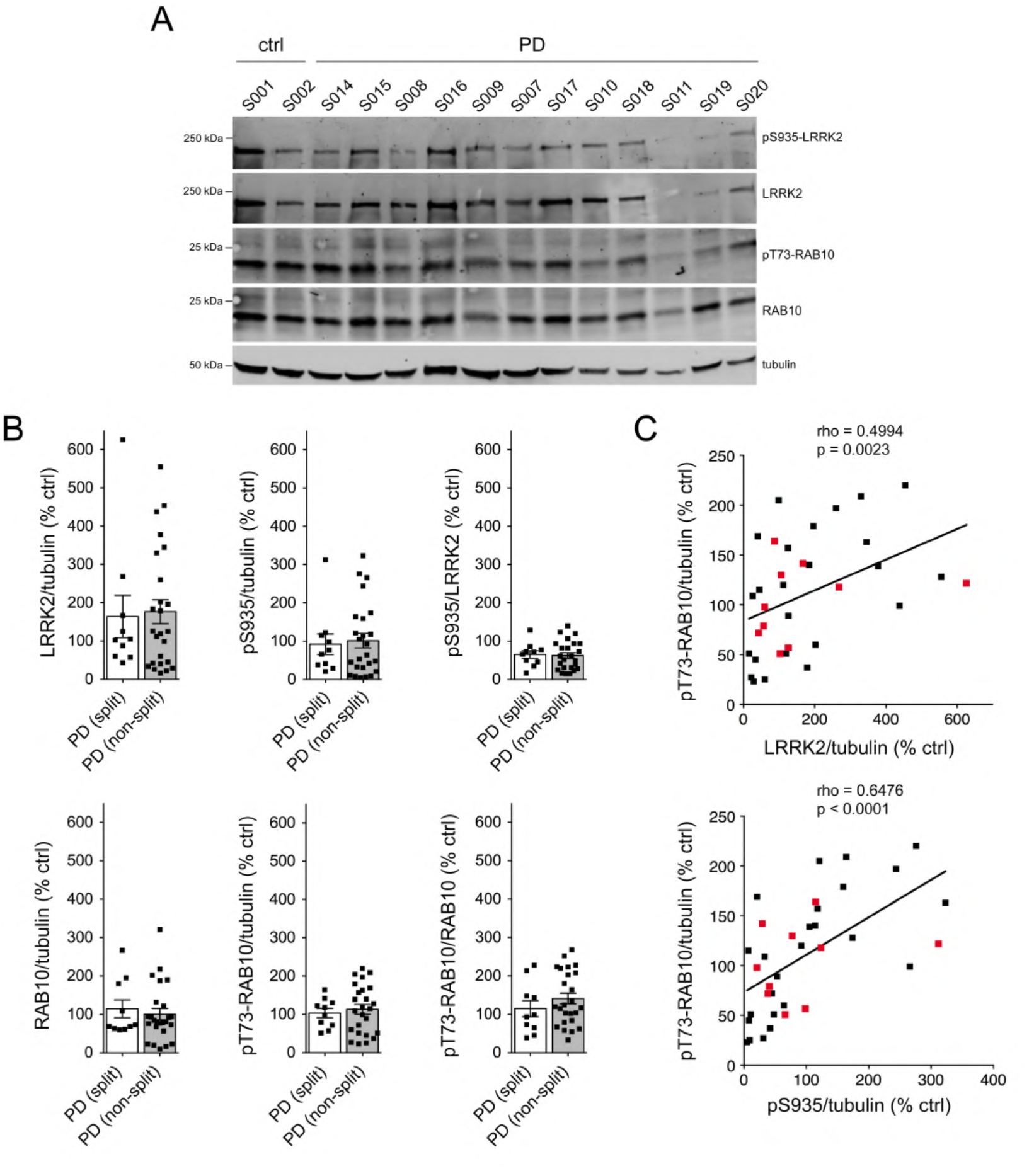
Analysis of LRRK2, S935-LRRK2, Rab10 and pT73-Rab10 levels in PD LCLs with or without a centrosome cohesion phenotype. (**A**) Example of two control and 12 PD LCL lines. Cells were lysed and extracts subjected to quantitative immunoblot analysis with the indicated antibodies and membranes were developed using Odyssey CLx scan Western Blot imaging system. pT73-Rab10 and total Rab10, as well as pS935-LRRK2 and total LRRK2 were multiplexed, and the same control line (GRA330) was run on every gel to compare samples run on different gels. (**B**) Immunoblots were quantified for LRRK2/tubulin, pS935/tubulin, pS935/LRRK2, Rab10/tubulin, pT73-Rab10/tubulin and pT73-Rab10/Rab10 as indicated, with no differences observed between PD LCL lines with or without a centrosome splitting phenotype. (**C**) Spearman correlation analysis between levels of LRRK2/tubulin and pT73-Rab10/tubulin (top) or pS935/tubulin and pT73-Rab10/tubulin (bottom). A significant association is observed between LRRK2 or S935-LRRK2 levels and pT73-Rab10 levels in PD LCLs. Red datapoints indicate the ten PD samples which display a centrosomal cohesion deficit. Rho and p values are indicated for each correlation analysis.

### Lysosomal damage causes LRRK2 kinase-mediated increases in pT73-Rab10 levels in both control and idiopathic PD patient-derived cells

Recent studies have shown that treatment of cells with the lysosome membrane-rupturing agent L-leucyl-L-leucine methyl ester (LLOMe) causes recruitment and activation of LRRK2 at damaged lysosomes which is associated with a potent increase in pT73-Rab10 levels (*53–55*). Consistent with these reports, LLOMe treatment caused a time-dependent increase in pT73-Rab10 levels which was reverted by MLi2 (**Fig. S1**). This correlated with an MLi2-sensitive accumulation of pT73-Rab10 near the centrosome (**Fig. S2**) and with a cohesion deficit which was reverted by MLi2 (**Fig. S3**). Moreover, the LLOMe-mediated alterations were observed in both healthy control and *G2019S-LRRK2* LCLs (**Fig. S1-3**). Hence, we reasoned that LLOMe-triggered LRRK2 activation may allow us to better detect potential differences in pT73-Rab10 levels amongst idiopathic PD LCL lines. Cells were treated with or without LLOMe and MLi2, and the LLOMe-induced increase in pT73-Rab10 levels determined for each cell line. LLOMe treatment induced a similar increase in pT73-Rab10 levels in control LCLs and in idiopathic PD LCLs irrespective of whether they displayed a centrosomal cohesion phenotype (**Fig. 3A,B**). The LLOMe-induced increase in pT73-Rab10 levels was reduced upon MLi2 treatment in most cases (**Fig. 3C, Figs. S4, S5**). Interestingly though, some idiopathic PD LCLs did not display a LLOMe-induced increase in pT73-Rab10 levels (**Fig. 3B**). Such lack of LLOMe-mediated potentiation of pT73-Rab10 levels marginally correlated with high basal levels of pT73-Rab10 in the absence of LLOMe treatment (**Fig. 3D**), and further work is required to determine whether these idiopathic PD patient-derived cells already harbor lysosomal damage and thus display maximal LRRK2 kinase activity. In either case, these data show that neither basal nor LLOMe-induced pT73-Rab10 levels correlate with the MLi2-sensitive centrosomal cohesion deficits observed in a subset of idiopathic PD LCLs.

**Figure 3.**
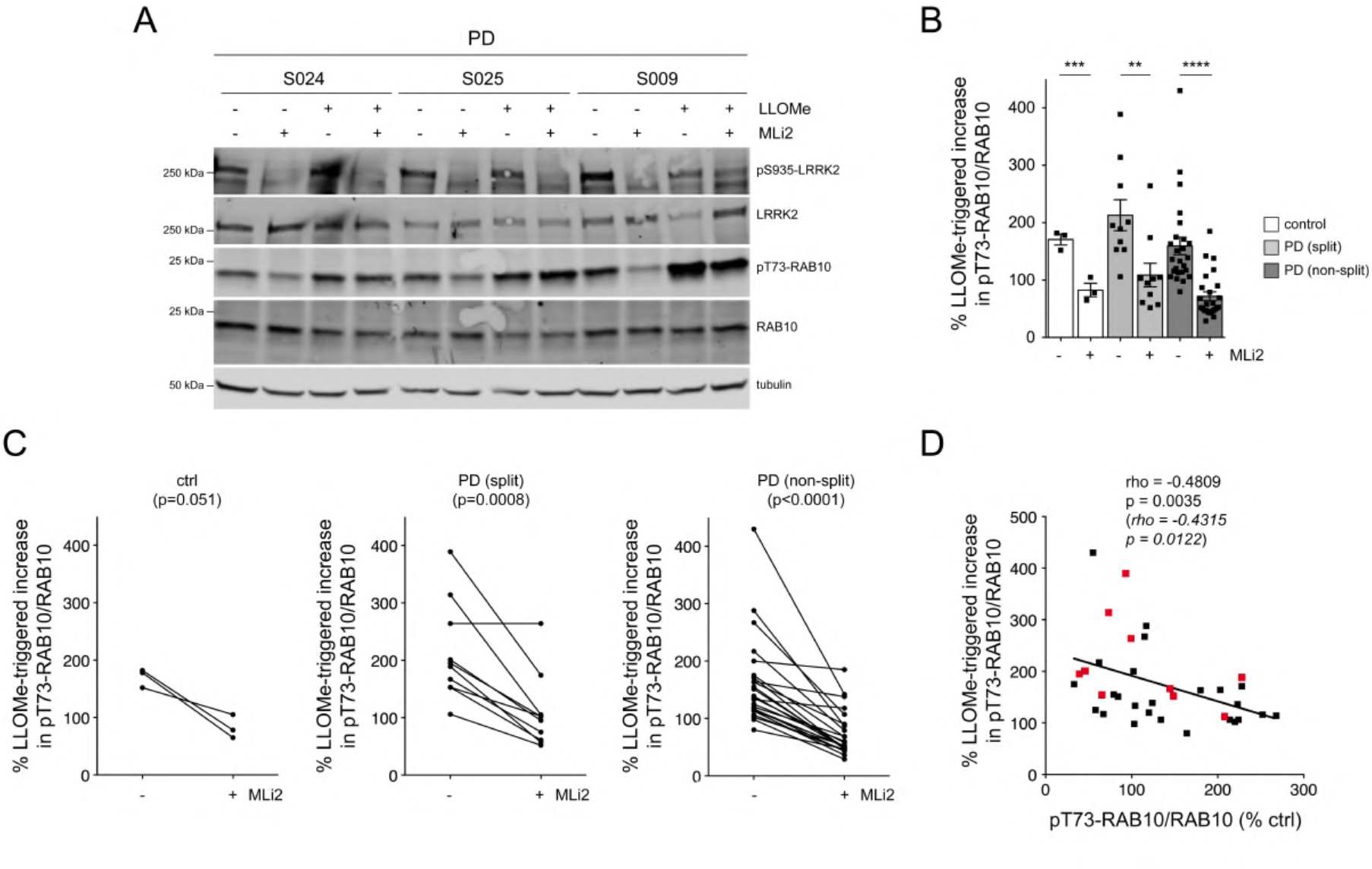
Effects of LLOMe treatment on LRRK2 kinase-mediated pT73-Rab10 levels. (**A**) Example of three PD LCL lines with or without treatment with LLOMe (1 mM) and MLi2 (50 nM) for 2 h as indicated. Cells were lysed and extracts subjected to multiplexed immunoblotting with the indicated antibodies. (**B**) The percentage of LLOMe-triggered increase in pT73-Rab10/Rab10 levels in the absence or presence of MLi2 was calculated for each LCL line. LLOMe triggers similar increases in pT73-Rab10/Rab10 levels in control and PD LCLs with or without a cohesion phenotype. Ctrl versus ctrl + MLi2 (p = 0.004); PD (split) versus PD (split) + MLi2 (p = 0.006); PD (non-split) versus PD (non-split) + MLi2 (p < 0.001): ****p < 0.001; ***p < 0.005; **p < 0.01. (**C**) Paired t-test analysis of LLOMe-triggered increase in pT73-Rab10/Rab10 levels from each cell line in the absence or presence of MLi2. Note that the LLOMe-triggered increase in pT73-Rab10/Rab10 levels is reduced by MLi2 treatment in most cell lines. (**D**) Spearman correlation analysis between the percentage of LLOMe-triggered increase in pT73-Rab10/Rab10 levels versus basal pT73-Rab10/Rab10 levels in the absence of LLOMe treatment. There is a negative correlation between basal pT73-Rab10/Rab10 levels and the efficacy of the LLOMe-mediated increase in pT73-Rab10/Rab10 levels. Red datapoints indicate the ten PD samples which display a centrosomal cohesion deficit. Rho and p values are indicated (in italics values without the two outliers).

### Identification of gene variants in idiopathic PD samples

We next wondered whether the centrosomal cohesion deficits in the idiopathic PD LCLs may be due to genetic alterations in select genes impacting upon centrosomal cohesion in a LRRK2 kinase activity-mediated manner. Whole exome sequencing revealed single nucleotide variants (SNVs) in PD-relevant genes for some LCL lines (**Table 2**). Amongst the idiopathic PD lines which displayed a centrosomal cohesion deficit, one line harboured a variant in the translational repressor *GIGYF2*, a gene at the *PARK11* locus with an unconfirmed link to PD (*56, 57*). Amongst the PD lines without a cohesion deficit, one displayed a variant in *ATP13A2*, and two displayed a known pathogenic missense mutation in *PRKN* (**Table 2**). Since heterozygous mutations in the *GBA* gene are the most frequent known genetic risk factor for PD, we additionally performed long-range PCR and Sanger sequencing of the *GBA* gene (*58*), which allowed for identification of the E326K variant known to be associated with PD risk in two lines without a cohesion phenotype (**Table 2**). Therefore, the MLi2-sensitive cohesion deficits observed in a subset of idiopathic PD samples are not due to mutations in *LRRK2* or in other genes related to PD risk, raising the possibility that variants unrelated to disease risk may be mediating the phenotype.

**Table 2.**
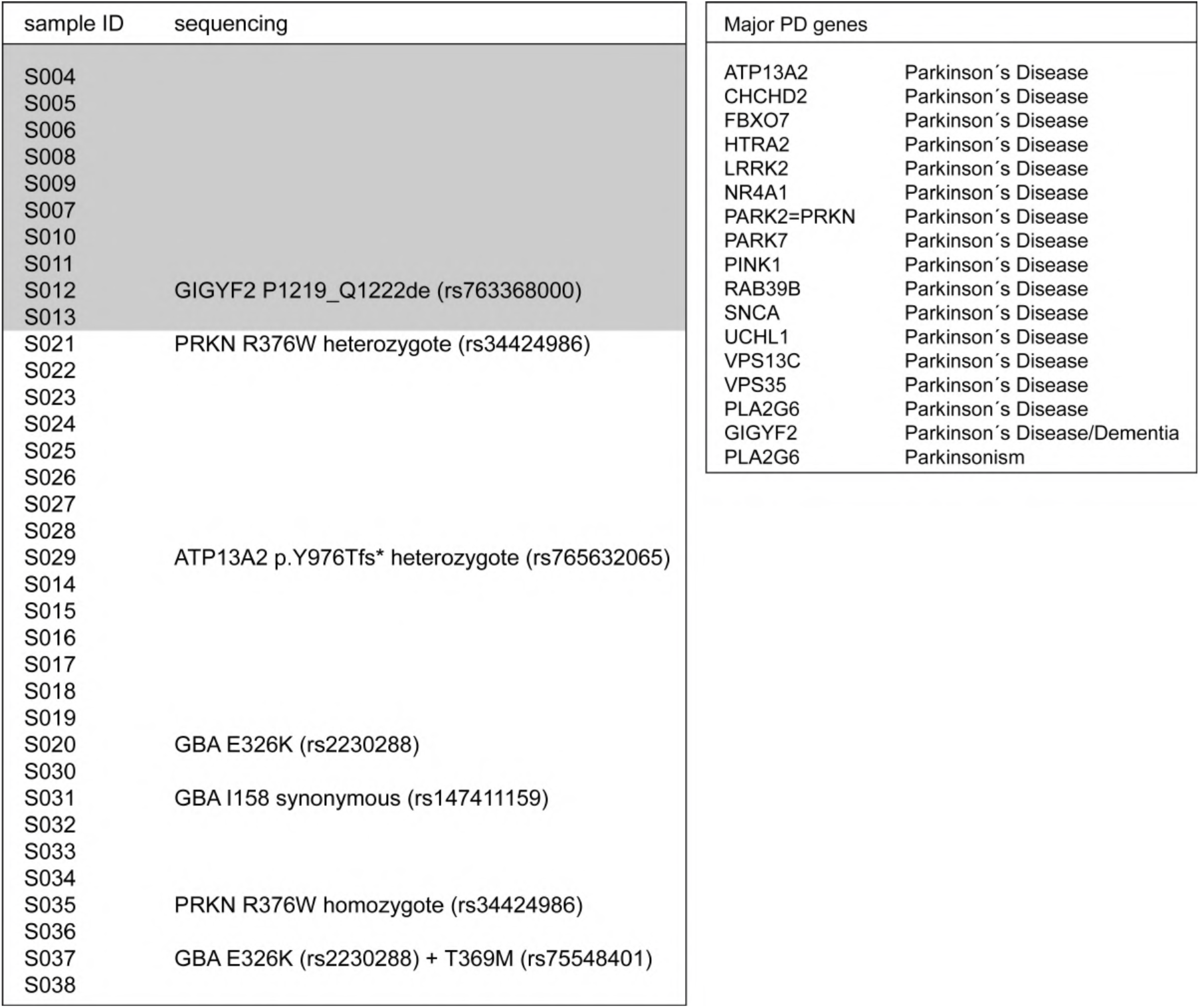
Several LCL lines display PD-relevant mutations. All 35 LCL PD lines were subjected to whole exome sequencing and long-range PCR and Sanger sequencing of the GBA gene, the most frequent known genetic risk factor for PD. The 10 PD LCL lines with a cohesion phenotype are boxed in light gray. Apart from GBA, the major PD genes analyzed are indicated to the right; de, deletion; fs, frameshift.

Whole exome sequencing data were next analysed to determine whether any other gene (or combination thereof) may be a better pharmacogenomic predictor than a PD gene mutation. Gene burden analyses indicated no significant burden of rare variants for any single gene after correcting for multiple comparisons (**Table S1, Table S2**). The highest association between the centrosomal cohesion phenotype and rare variants was found within the *TBC1D3D* gene (**Table 3**), a member of the TBC1D3 family which may act as an effector protein for Rab5 (*59*) (p = 1.44 x 10-5). Another association was observed with rare variants in *NOTCH2NLC* (p = 0.00055851) (**Table 3**), and repeat expansions in *NOTCH2NLC* have recently been detected in idiopathic PD cases (*60, 61*). Pathway analysis indicated a significant overrepresentation of a KEGG pathway (hsa04612: Antigen processing and presentation, FDR = 0.0032, enrichment ratio 27.71) comprising the *HLA-DPB1*, *KIR2DL4*, *KIR2DS4* and *KIRDL1* genes (**Table S1; Table S2**). The same gene list was also significantly enriched for a specific domain (PF06758: Repeat of unknown function (DUF1220), FDR = 1.14 e-07) and comprising the *NBPF10*, *NBPF12*, *NBPF14*, *NBPF9* and *PDE4DIP* genes (**Table 3**).

**Table 3.**
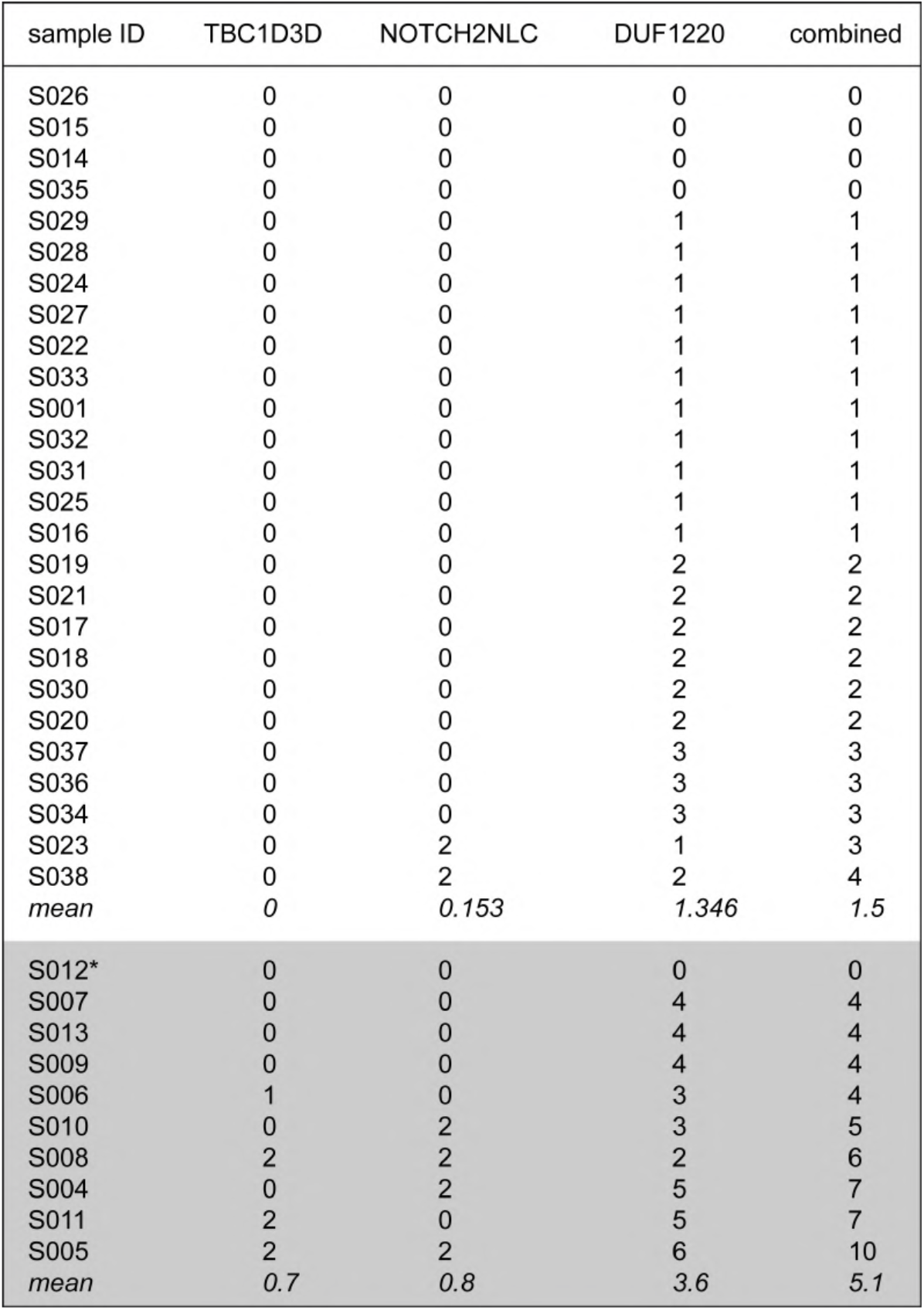
Gene burden analysis for rare variants employing whole exome sequencing data from the different PD LCL lines. Whilst there is no significat burden of rare variants which correlates with the centrosomal cohesion phenotype for any single gene after correcting for multiple comparisons, there is an association between the centrosome phenotype and rare variants within the *TBC1D3D* gene and the *NOTCH2NLC* gene. Pathway analysis also indicates enrichment for a list of genes enriched for the DUF1220 domain. Numbers indicate the number of rare variants identified for each LCL line. The 10 PD LCL lines with a cohesion phenotype are boxed in light gray. Note that S012 is the line containing a mutation in *GIGYF2*.

Interestingly, *TBC1D3*, *NOTCH2NL* and *DUF1220* domain-containing genes are all hominoid-specific genes which have undergone duplications during evolution, with *NOTCH2NL/DUF1220* duplicating as a two-gene module, followed by further hyperamplification of DUF1220 domains encoded by the *NBPF* genes (*62, 63*). Expression of either TBC1D3, NOTCH2NL or DUF1220 protein domains drives proliferation of neural stem and progenitor cells and promotes cortical brain expansion and folding (*64–67*). Whilst further validation using *in vitro* cellular models and knockdown or overexpression studies are warranted, these findings are consistent with a role for these gene products in centrosome-mediated effects which may affect a celĺs response to MLi2.

### Cohesion deficits in LCLs from distinct *LRRK2* mutation carriers

It is unknown whether centrosomal deficits are a phenotype common to carriers of distinct *LRRK2* mutations. To address this question, we analyzed another cohort of subjects (n=10 controls, n=12 *R1441G-LRRK2* PD, n=9 *R1441G-LRRK2* non-manifesting carriers (NMC), n=7 *G2019S-LRRK2* PD, n=6 *G2019S-LRRK2* NMC, n=4 idiopathic PD) (**Table 4**). Several tubes of highly concentrated peripheral blood mononuclear cells (PBMCs) were obtained from all patients, and 1-2 tubes were employed to generate EBV-transformed LCLs.

**Table 4.**
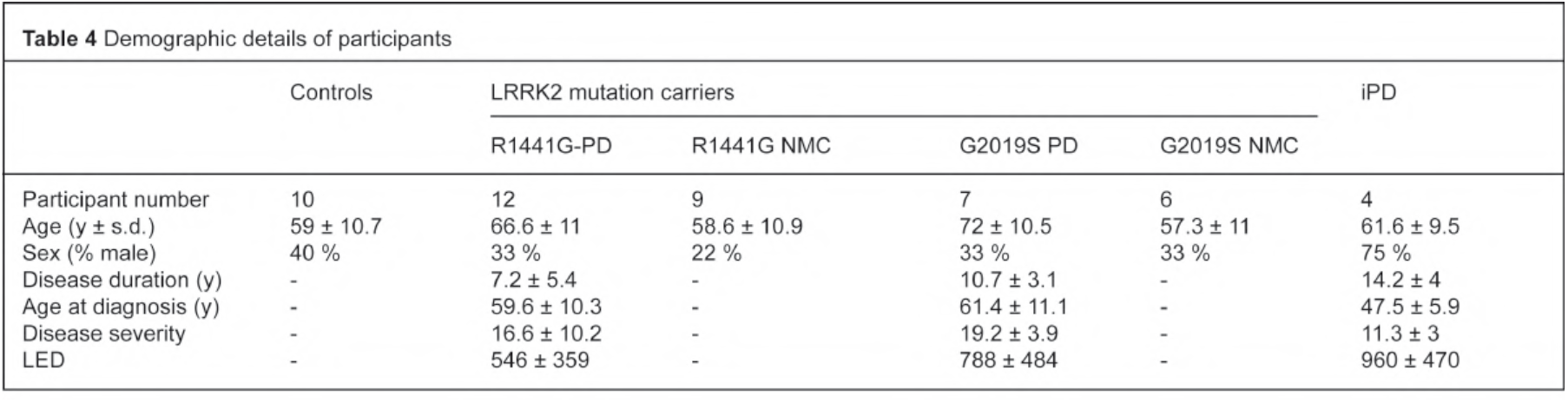
Demographic details for PD and control patients. Data shown are mean ± s.d. Disease severity was measured using the Movement Disorders Society Unified Parkinso’s Disease Rating Scale (MDS-UPDRS) part III, and LED is the calculated L-dopa-equivalent dose. All patients were sequenced for the G2019S and R1441G LRRK2 mutations. NMC, non-manifesting carriers.

To increase the ease and speed of analysis, cells were only stained with an antibody against pericentrin. Based on a frequency histogram of the distances between two pericentrin-positive dots (*68*) from a control LCL line, the mean distance was around 1.2 μm (**Fig. S6**), which is smaller than that previously determined when analyzing dots positive for both pericentrin and γ-tubulin (*52*). In addition, around 60% of cells displayed two pericentrin-positive dots (**Fig. S6**), which is three times higher than the amount of LCLs in G2/M phase as determined by flow cytometry (*52*). Bromodeoxyuridine (BrdU) incorporation assays to detect DNA synthesis which occurs concomitant with centrosome duplication events further indicated only a small percentage of LCLs in S phase of the cell cycle (**Fig. S6**). Therefore, analysis of the distance between two pericentrin-positive dots not only measures the cohesion between duplicated centrosomes from cells in S and G2 phases, but also the cohesion between centrioles from cells in G1 phase of the cell cycle. Such centrosome/centriole (C/C) splitting was defined as the percentage of cells with a distance between two pericentrin-positive dots greater than 1.3 μm (**Fig. S6**).

Analysis of C/C cohesion revealed a deficit in both *R1441G-LRRK2* and *G2019S-LRRK2* PD LCLs which was reverted by MLi2 in all cases (**Fig. 4A-C**). The C/C cohesion deficit was reflected by an overall increase in the mean distance between the two pericentrin-positive structures (**Fig. S6**) and was not associated with changes in the percentage of cells displaying two dots (**Fig. S7**). A significant C/C splitting deficit was also observed in the four idiopathic PD LCLs (**Fig. 4B,C**). Importantly, both *R1441G-LRRK2* and *G2019S-LRRK2* mutation NMCs displayed a C/C cohesion deficit relative to healthy age-matched controls which was sensitive to MLi2 in most cases (**Fig. 4B,C**). C/C splitting in *R1441G-LRRK2* PD and *G2019S-LRRK2* PD strongly predicted a PD diagnosis with a ROC area under the curve c-statistic of 1.0, with a lower value obtained for *R1441G-LRRK2* and *G2019S-LRRK2* NMCs (**Fig. S8**). No significant correlations were found between C/C splitting levels and any demographic or clinical data (not shown).

**Figure 4.**
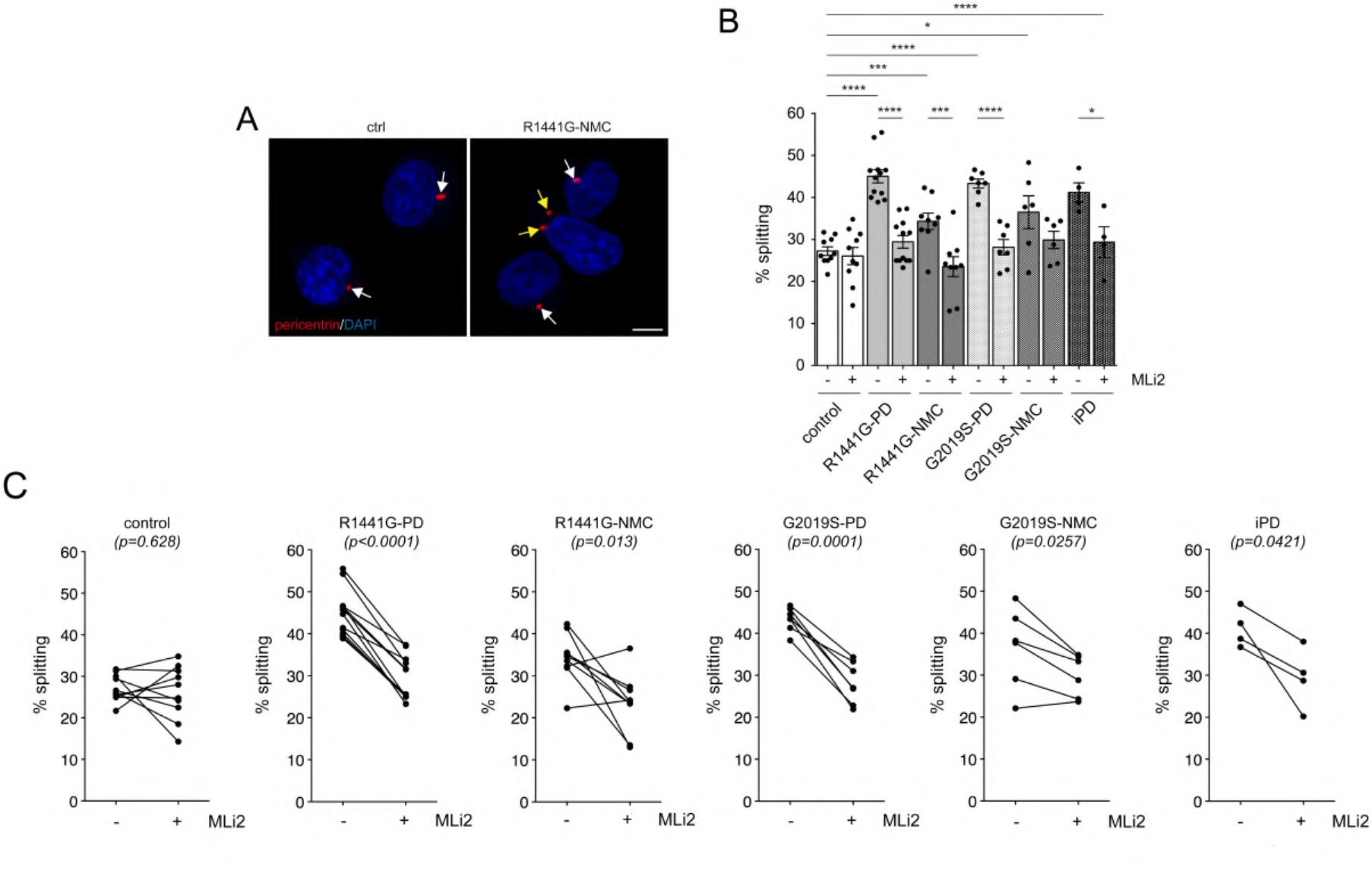
C/C cohesion deficits in *R1441G-LRRK2* and *G2019S-LRRK2* LCLs. (**A**) Example of a healthy control (ctrl) and an *R1441G-LRRK2* NMC LCL line stained for the centrosomal marker pericentrin and DAPI. Arrows point to pericentrin-positive dots, yellow arrows to duplicated split pericentrin-positive dots. Scale bar, 5 μm. (**B**) The cohesion phenotype was quantified from 150-200 cells per line from 10 control, 12 *R1441G-LRRK2* PD, 9 *R1441G-LRRK2* NMC, 7 *G2019S-LRRK2* PD, 6 *G2019S-LRRK2* NMC and 4 idiopathic PD patient LCLs in either the absence or presence of MLi2 (50 nM, 2 h) as indicated. Ctrl versus R1441G mutation (p < 0.0001); ctrl versus R1441G NMC (p = 0.0039); ctrl versus G2019S mutation (p < 0.0001); ctrl versus G2019S NMC (p = 0.012); ctrl versus idiopathic PD (p < 0.0001); R1441G mutation versus R1441G mutation + MLi2 (p < 0.0001); R1441G NMC versus R1441G NMC + MLi2 (p = 0.002); G2019S mutation versus G2019S mutation + MLi2 (p < 0.0001); idiopathic PD versus idiopathic PD + MLi2 (p = 0.033). ****p < 0.001; ***p < 0.005; **p < 0.01; *p < 0.05. (**C**) Paired t-test analysis of cohesion deficits from each cell line in the absence or presence of MLi2 as indicated.

Quantitative immunoblotting showed highly variable levels of total LRRK2 which were not significantly different amongst control, *R1441G-LRRK2* PD, *R1441G-LRRK2* NMC, *G2019S-LRRK2* PD, *G2019S-LRRK2* NMC and idiopathic PD LCLs, whilst the levels of total Rab10 were more similar (**Fig. 5A,B**). There were no significant differences in pT73-Rab10 levels amongst control, *LRRK2* mutation PD, *LRRK2* NMC and idiopathic PD samples, even though pT73-Rab10 levels were susceptible to MLi2 treatment in most cases (**Fig. 5C, Fig. S9, Fig. S10**). Correlation analysis indicated a significant positive correlation between LRRK2 levels and pT73-Rab10 levels (**Fig. 5D**), but there was no correlation between the extent of C/C splitting and either pT73-Rab10 or total LRRK2 levels (**Fig. S11**). Hence, LRRK2 kinase activity-mediated C/C cohesion deficits are present in both manifesting and non-manifesting *LRRK2* mutation carriers but do not correlate with increased pT73-Rab10 or LRRK2 levels as assessed by quantitative Western blotting techniques. Since the C/C splitting phenotype results from the pericentrosomal accumulation of pT73-Rab10, we stained a subset of healthy control, *R1441G-LRRK2* PD and *R1441G-LRRK2* NMC LCLs for both pericentrin and pT73-Rab10 (**Fig. S12**). However, and similar to what we observed with Western blotting techniques, the percentage of LCLs displaying detectable pT73-Rab10 accumulation was highly variable and was not different between control and *R1441G-LRRK2* mutation carriers (**Fig. S12**).

**Figure 5.**
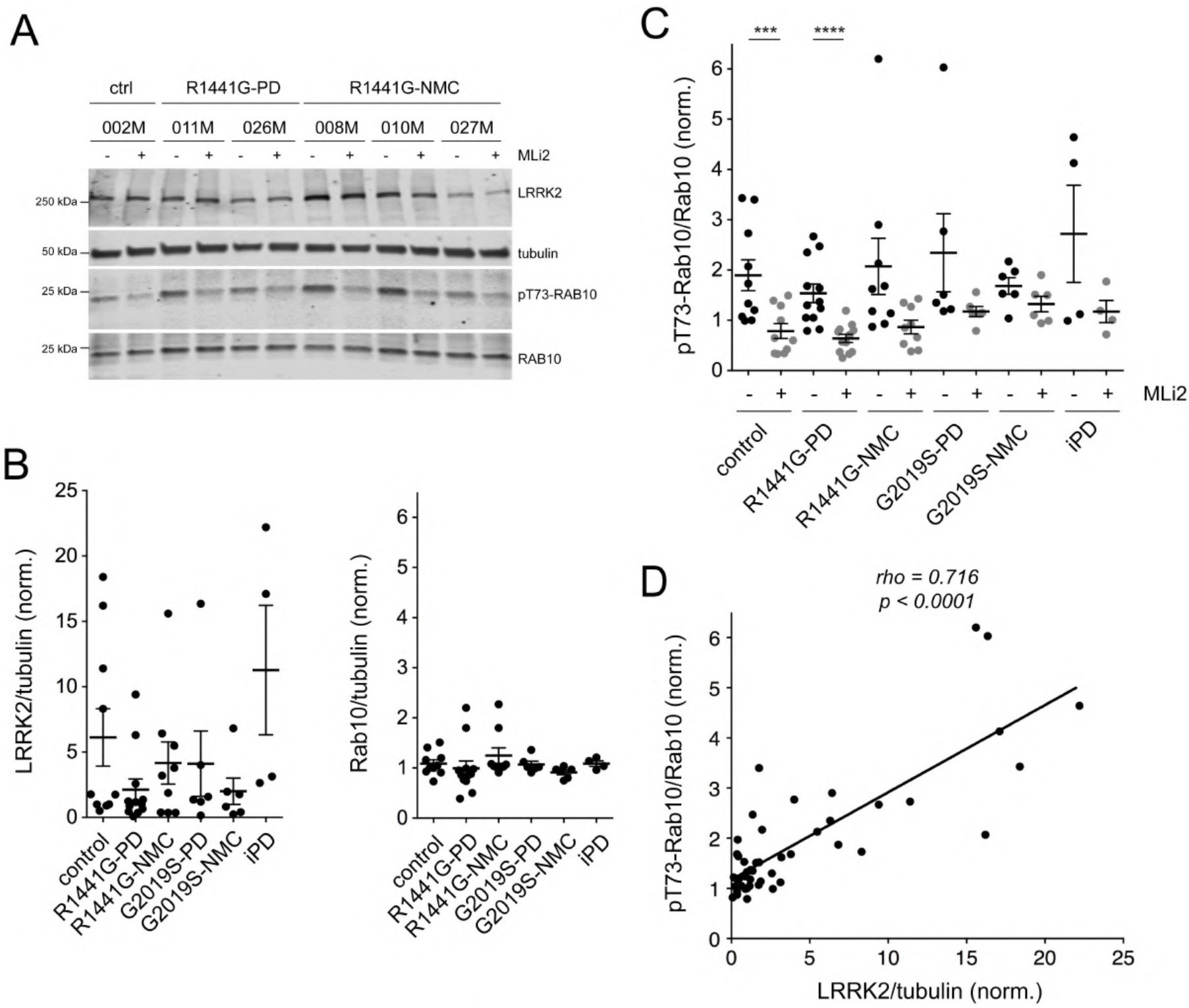
Analysis of LRRK2, Rab10 and pT73-Rab10 levels in *R1441G-LRRK2* and *G2019S-LRRK2* LCLs. (**A**) Example of control, *R1441G-LRRK2* PD and *R1441G-LRRK2* NMC LCL lines in the absence or presence of MLi2 (50 nM, 2 h) as indicated. Cell extracts were subjected to multiplexed quantitative immunoblot analysis with the indicated antibodies, and membranes were developed using Odyssey CLx scan Western Blot imaging system. The same control line (002M) was run on every gel as an internal standard to compare samples run on different gels. (**B**) Immunoblots were quantified for LRRK2/tubulin levels (left) and Rab10/tubulin levels (right). Note large variability in the total LRRK2 levels amongst different LCL lines. (**C**) Immunoblots were quantified for pT73-Rab10/Rab10 levels, with no significant differences observed between control and PD LCL lines. Ctrl versus ctrl + MLi2 (p = 0.004); *R1441G-LRRK2* PD versus *R1441G-LRRK2* PD + MLi2 (p = 0.0002). ****p < 0.001; ***p < 0.005. (**D**) Spearman correlation analysis between levels of pT73-Rab10/Rab10 and LRRK2/tubulin from all LCL lines analyzed. Rho and p values are indicated. A significant association is observed between the total levels of LRRK2/tubulin and the levels of pT73-Rab10/Rab10.

### Cohesion deficits in PBMC lymphocytes from *LRRK2* mutation carriers

PBMCs are routinely used for biomarker studies and are currently employed in the clinic to measure target engagement for LRRK2 and other PD-relevant therapeutics. Since the majority (70-90%) of human PBMCs is composed of lymphocytes (*69*), we evaluated whether the C/C cohesion phenotype was also observed in non-immortalized blood-derived peripheral cells. Staining of control and *G2019S-LRRK2* PD PBMC preparations obtained from the LRRK2 Biobanking Initiative revealed extensive cell death and debris, even though lymphocytes could be identified by their known small cell size with a nuclear diameter < 10 μm (**Fig. S13**). In contrast, monocytes displayed a larger and typical kidney bean-shaped nucleus which was independently confirmed by employing purified monocyte preparations (**Fig. S13**).

Our PBMC preparations had been collected and cryopreserved under conditions optimal for subsequent cell biological analysis, and they were next employed to determine the C/C cohesion phenotype. Analysis of cells with a nuclear diameter < 10 μm indicated a mean distance between two pericentrin-positive dots of around 1.1 μm, with around 60% of cells displaying a two-dot phenotype (**Fig. S14**). BrdU incorporation assays showed a neglible amount of cells in S phase (**Fig. S6**), again consistent with the notion that we are also detecting centriolar cohesion deficits in G1 phase of the cell cycle. C/C splitting was defined when the distance was > 1.3 μm, and a test-retest reliability assay employing another cryopreserved PBMC tube from the same healthy control patient indicated that the phenotype was stable (**Fig. S14**).

C/C cohesion was analyzed from lymphocytes with a nuclear diameter < 10 μm from all patient-derived PBMCs. As compared to healthy age-matched controls, *R1441G-LRRK2* PD and *G2019S-LRRK2* PD lymphocytes displayed a C/C splitting phenotype which was reverted by MLi2 in all cases (**Fig. 6A-C**). The cohesion phenotype was reflected by an overall increase in the mean distance between the pericentrin-positive dots without changes in the percentage of cells displaying two dots (**Fig. S15**). An MLi2-sensitive splitting phenotype was also observed in lymphocytes from idiopathic PD patients and in lymphocytes from *R1441G-LRRK2* and *G2019S-LRRK2* NMCs (**Fig. 6B,C, Fig. S8**). Finally, over the entire sample cohort there was a good correlation between the C/C cohesion phenotype as determined from PBMCs versus that determined from LCLs (**Fig. S16**). Altogether, these data indicate that C/C cohesion deficits are a blood-based cellular biomarker which is detectable in lymphocytes from *LRRK2* mutation carriers and some idiopathic PD patients as compared to healthy controls and which is responsive to LRRK2 kinase inhibition.

**Figure 6.**
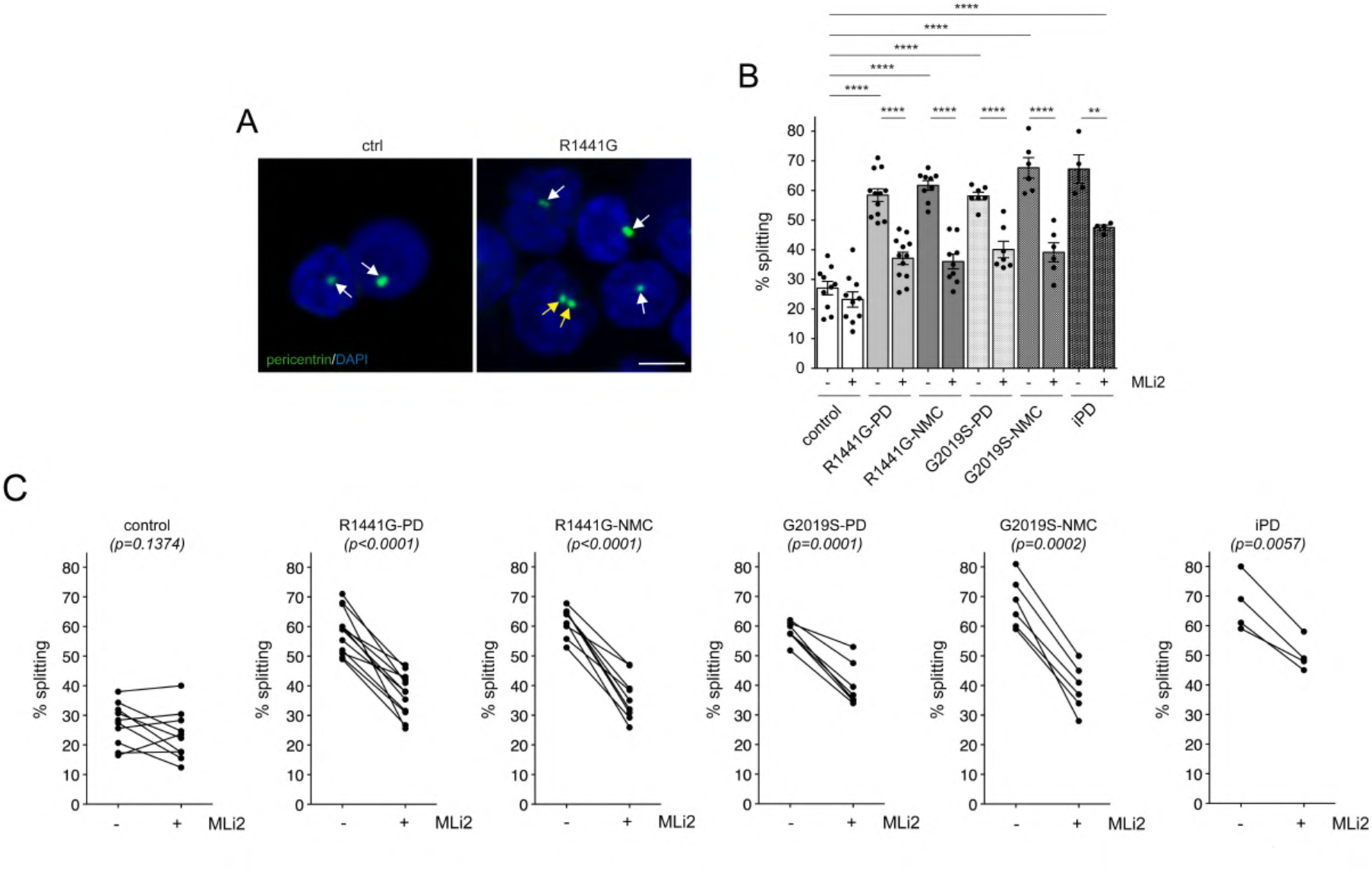
C/C cohesion deficits in lymphocytes from *R1441G-LRRK2* and *G2019S-LRRK2* mutation carriers. (**A**) Example of healthy control (ctrl) and *R1441G-LRRK2* PD lymphocytes from PBMC preparations stained for pericentrin and DAPI. Arrows point to pericentrin-positive structures, yellow arrows to two pericentrin-positive structures displaying a split phenotype. Scale bar, 5 μm. (**B**) The C/C splitting phenotype was quantified from 150-200 cells from 10 control, 12 *R1441G-LRRK2* PD, 9 *R1441G-LRRK2* NMC, 7 *G2019S-LRRK2* PD, 6 *G2019S-LRRK2* NMC and 4 idiopathic PD patients in either the absence or presence of MLi2 (200 nM, 30 min) as indicated. Ctrl versus R1441G mutation (p < 0.0001); ctrl versus R1441G NMC (p < 0.0001); ctrl versus G2019S mutation (p < 0.0001); ctrl vrsus G2019S NMC (p < 0.0001); ctrl versus idiopathic PD (p < 0.0001); R1441G mutation versus R1441G mutation + MLi2 (p < 0.0001); R1441G NMC versus R1441G NMC + MLi2 (p < 0.0001); G2019S mutation versus G2019S mutation + MLi2 (p < 0.0001); G2019S NMC versus G2019S NMC + MLi2 (p = 0.0001); idiopathic PD versus idiopathic PD + MLi2 (p = 0.006). ****p < 0.001; **p < 0.01. (**C**) Paired t-test analysis of C/C cohesion deficits from each cell line in the absence or presence of MLi2 as indicated.

## Discussion

Here, we probed for LRRK2 kinase-mediated centosomal alterations in two different cohorts of PD patient-derived cells. Based on whole exome sequencing and long-range PCR and Sanger sequencing of idiopathic PD patients, none of the patient samples with a cohesion deficit had mutations in genes known to cause PD. Cohesion deficits were observed in a significant percentage of idiopathic PD patients and were reverted by a LRRK2 kinase inhibitor in all cases. These data are consistent with previous reports (*12, 31*) and indicate that the wildtype LRRK2 kinase activity is increased in at least a subset of idiopathic PD patients.

All patients with PD due to either the *R1441G* or the *G2019S* mutation in *LRRK2* and all non-manifesting *LRRK2* mutation carriers showed an MLi2-reversible cohesion deficit. These data demonstrate that cohesion deficits serve as a robust cellular biomarker for the presence of pathogenic LRRK2 in peripheral blood-derived cells which may begin to appear prior to a clinical PD diagnosis. In contrast, *LRRK2* PD or idiopathic PD samples did not display an increase in the levels of pT73-Rab10 as compared to age-matched healthy controls. These findings are consistent with previous studies which also failed to detect a robust increase in pT73-Rab10 levels in *LRRK2* PD or idiopathic PD patients (*20–24, 26–28, 52*). It is known that total LRRK2 levels in PBMCs and LCLs vary widely among individuals (*24, 52*), which may mask an overall increase in pT73-Rab10 levels. We also observed vastly different levels of total LRRK2, and a positive correlation between total LRRK2 levels and pT73-Rab10 levels. However, the cohesion deficits did not correlate with the levels of total LRRK2 or pT73-Rab10. Since the cohesion deficits were reverted by short-term treatment with MLi2 in all cases, they are driven by a LRRK2 kinase-mediated process, perhaps influenced by additional cellular effects of LRRK2 and involving additional kinase substrates. In either case, these data indicate that pT73-Rab10 levels accurately reflect target engagement of LRRK2 kinase inhibitors but are not able to stratify PD patients with increased LRRK2 activity.

When treating idiopathic PD LCLs with LLOMe to trigger lysosomal damage-induced LRRK2 activation, we observed that some samples did not display a LLOMe-induced increase in pT73-Rab10 levels. These included the two patient samples with mutations in *PRKN* as well as the two patient samples with mutations in *GBA*. The latter is consistent with the idea that lysosomal deficits cause activation of wildtype LRRK2. In addition, these data suggest that quantifying LLOMe-mediated alterations in pT73-Rab10, rather than pT73-Rab10 levels *per se*, may be a suitable approach to nominate PD patients with lysosomal dysfunction and consequent LRRK2 activation. More extensive studies with LCLs harbouring distinct *GBA* mutations are required to corroborate these observations.

A gene burden analysis of the idiopathic PD samples did not reveal a significant burden of rare variants for any single gene which correlated with the cohesion phenotype. Whilst the PD cohort available here was statistically underpowered for such analysis, it nevertheless highlighted several candidate modifier genes which may impact upon the cohesion phenotype. Interestingly, rare variants which associated with the cohesion phenotype were detected in three groups of genes (*TBC1D3*, *NOTCH2NL*, DUF1220 domain-containing genes) which are all hominoid-specific, implicated in the proliferation of neural stem and progenitor cells and involved in cortical brain expansion and folding (*62–67*). Of note, one of the DUF1220 domain-containing genes is *PDE4DIP* (phosphodiesterase 4D-interacting protein, also called myomegalin). Myomegalin is a paralogue of CDK5RAP2 which localizes to the centrosome and recruits the cyclic nucleotide phosphodiesterase PDE4D to this location (*70, 71*). Myomegalin loss dislocates PDE4D from the centrosome causing local PKA overactivation and inhibition of hedgehog (Hh) signaling, which is followed by decreased neural precursor cell proliferation (*72*). We have previously shown that the cohesion deficits mediated by mutant LRRK2 are due to the kinase activity-mediated displacement of centrosomal CDK5RAP2 (*42*). In future work, it will be important to determine whether and how the myomegalin/PDE4D pathway impacts upon cohesion in a manner mitigated by LRRK2 kinase inhibition.

The MLi2-sensitive centrosomal defects as quantified here reflect defects in the cohesion of duplicated centrosomes as observed in S and G2 phases of the cell cycle, but also defects in the cohesion of centrioles as observed in G1 phase. Both centrosome and centriole cohesion are mediated by a small set of proteins including CDK5RAP2 (*45, 46*) and can be monitored by staining for pericentrin, a marker for the pericentriolar material (*73*). Pericentrin has also been shown to interact with kinesin-1 to drive centriole motility (*49*). Given the reported links between Rab phosphorylation, motor adaptor protein recruitment and microtubule-mediated transport processes (*53, 74, 75*), it will be interesting to determine whether the LRRK2-mediated cohesion deficits are due to a similar phospho-Rab/motor protein interaction which is followed by inappropriate microtubule-mediated transport. In either case, and given that peripheral blood-derived cells including monocytes, neutrophils and lymphocytes are non-ciliated cells (*76*), the LRRK2-mediated centrosomal defects may be particularly pronounced in those cells and thus able to serve as robust cellular biomarker for PD due to increased LRRK2 activity.

Recent studies indicate that mitocondrial DNA damage may be another potential biomarker for PD which is reversed by LRRK2 kinase inhibitors and detectable in LCLs and PBMCs, even though it does not correlate with increased pT73-Rab10 levels (*77–80*). It will be important to determine whether idiopathic PD patients with a cohesion deficit also display high levels of mitochondrial DNA damage, or whether these biomarkers are detecting distinct cellular outcomes of increased LRRK2 activity. Similarly, it will be interesting to determine whether idiopathic PD patients with pre-existing lysosomal damage as assessed by the lack of a LLOMe-stimulated increase in pT73-Rab10 levels display high levels of mitochondrial DNA damage. Importantly, cohesion deficits as determined in this study are not only observable in immortalized lymphocytes but also in PBMCs, and thus easily translatable to clinical settings.

A main limitation of the present study is the need for high-resolution confocal imaging. Future studies employing flow cytometry-based imaging and automated image quantification may allow for higher-throughput determination of cohesion deficits in peripheral blood from PD patients. We have shown cohesion deficits in *LRRK2* mutation carriers and in some early-stage idiopathic PD patients which were reversed by MLi2 in all cases. Whilst these results indicate that the cohesion deficits reflect a LRRK2-mediated process, it is unknown how they may be modulated by disease progression or disease severity. In addition, it is unknown whether cohesion defects as determined in the blood are mirrored by centrosomal/ciliary defects in the brain (*40, 44*), and evidence for LRRK2 kinase-mediated ciliary defects in the human brain is currently lacking. It is also unknown whether the cohesion defects observed in lymphocytes trigger peripheral immune responses which then initiate a neurodegenerative cascade (*81*), possibly in the absence of centrosomal/ciliary defects in the brain. Finally, another limitation of our study is sample size, and our present findings require replication in a larger patient cohort.

In sum, we here provide evidence that cohesion deficits are present in primary lymphocytes from *LRRK2* PD patients and non-manifesting *LRRK2* mutation carriers. They are also observed in a subset of early-stage idiopathic PD patients and are sensitive to LRRK2 kinase treatment. These data provide strong evidence for LRRK2-mediated cohesion deficits in a subset of idiopathic PD patients and support the future inclusion of this readout as a blood-based biomarker for patient enrichment in clinical trials with LRRK2-related therapeutics.

## Materials and Methods

### Study participants

For the first cohort, subjects were recruited with written informed consent and the study was approved by the University of New South Wales human research ethics committee. Patients were examined by neurologists specialized in movement disorders. None of the controls had a family history of PD.

For the second cohort, subjects were recruited at the Donostia University Hospital in San Sebastian with written informed consent, and the study was approved by the local ethics committee. Patients were examined by neurologists specialized in movement disorders, and the Movement Disorders Society Unified Parkinso’s Disease Rating Scale (MDS-UPDRS) part III was used to define motor symptom severity. L-dopa-equivalent dose (LED) was calculated for all patients. Subjects participating in the study donated blood samples for DNA extraction and routine LRRK2 genotyping, and peripheral blood mononuclear cells (PBMCs) for direct analysis as well as for lymphocyte immortalization. None of the controls had a family history of PD, and all control and idiopathic PD samples were negative for the *G2019S-LRRK2* or *R1441G-LRRK2* mutation.

### Peripheral blood mononuclear cell (PBMC) isolation and transformation

For the San Sebastian cohort, 35 ml of patient-derived blood was subjected to immediate purification of PBMCs using BD Vacutainer (CPT) Sodium Heparin tubes, and purified PBMCs were frozen at high cell density (1 x 10^7^ cells/tube) in cryopreservation medium (90% FBS, 10% DMSO). For all patients, 1-2 cryovials of purified PBMCs were employed to generate LCLs, and the remainder of cryovials was employed for direct analysis as described below. Lymphocytes were immortalized with Epstein-Barr virus (EBV) according to standard transformation protocols (*82*) which include cell separation by gradient centrifugation and lymphocyte growth enhancement with 1% (v/v) of the mitogenic phytohemagglutinin-M (PHA-M, ThermoFisher 10576015).

Samples (n=10) collected through the LRRK2 Biobanking Initiative at Columbia University were used for assay development. PBMCs were collected from patients and control subjects at the Movement Disorder Division in the Department of Neurology at Columbia University Irving Medical Center. Participants were screened for the LRRK2 G2019S mutation as well as several GBA1 mutations and variants (*28*). All participants provided written informed consent to take part in the study, and the study protocol was approved by the IRBs of both Columbia Univers0ity (CUIMC). PBMCs were isolated using standard protocols as previously described (*83*). Briefly, BD Vacutainer tubes (Sodium Heparin or Sodium Citrate) were used to collect whole blood from study participants. Blood was diluted 1 x in sterile PBS and transferred to Ficoll-containing Leucosep tubes (Griener) and centrifuged at 1000 x g for 10 min at room temperature. The upper plasma phase was then extracted and centrifuged at 300 x g for 15 min at room temperature. The banded cells were collected and washed once in PBS and centrifuged again at 300 x g for 10 min. Cells were resuspended in RPMI medium containing 40% FBS and 10% DMSO, counted and aliquoted at 3 x 10^6^ viable cells per cryovial. Frozen cells were stored at -80 °C.

### Cell culture and treatments

LCLs were grown as previously described (*52*). Briefly, cells were maintained in RPMI 1640 medium (ThermoFisher, 21870076) supplemented with 20% fetal bovine serum (ThermoFisher, 10437028), 2% L-glutamine (ThermoFisher, 25030081), 20 units/ml penicillin and 20 μg/ml streptomycin (ThermoFisher, 15140122) in T75 flasks (ThermoFisher, 156499) in 5% CO_2_ at 37 °C. Cells were maintained at a density of 10^6^ cells/ml, with cell density monitored every other day using trypan blue staining. Cell clumps were dispersed by pipetting, and 500’000 cells/ml treated in 1.5 ml tubes with DMSO or 50 nM MLi2 for 2 h before processing for immunocytochemistry. HEK293T and A549 cells were cultured as previously described (*42*).

Cryopreserved PBMCs were quickly thawed in a 37 °C waterbath, and transferred to 50 ml tubes containing 10 ml prelaid warm growth medium (RPMI 1640 medium with 20% fetal bovine serum, 2% L-glutamine, 20 units/ml penicillin and 20 μg/ml streptomycin), and cells centrifuged at 300 x g for 5 min at room temperature. The cell pellet was gently resuspended, and cells treated in growth medium in 12-well plates (1 x 10^6^ cells/well) with either DMSO or MLi2 (200 nM) for 30 min before processing for immunocytochemistry as described below.

### Immunocytochemistry

Coverslips (13 mm diameter) were placed into 24-well plates and coated with Cell-Tak and Tissue Adhesive solution (Corning, 354240) according to manufactureŕs instructions. After 30 min incubation at 37 °C, the solution was removed and coverslips were rinsed twice with distilled water followed by air-drying. LCLs or PBMCs (500’000 cells/coverslip) were added to dry coated coverslips and attached by slight centrifugation at 20 x g for 5 min at room temperature (without brake).

LCLs and PBMCs were fixed with 2% paraformaldehyde (PFA) in PBS for 20 min at room temperature followed by 5 min of ice-cold methanol fixation (for γ-tubulin staining only). Upon fixation, cells were permeabilized with 0.2% Triton-X100/PBS for 10 min at room temperature and blocked for 1 h in 0.5% BSA (w/v) (Millipore, 126579) in 0.2% Triton-X100/PBS (blocking buffer). Coverslips were incubated with primary antibodies in blocking buffer at 4 °C overnight. The following day, coverslips were washed three times for 10 min in 0.2% Triton-X100/PBS, followed by incubation with secondary antibodies in 0.2% Triton-X100/PBS for 1 h at room temperature. Coverslips were washed three times in 0.2% Triton-X100/PBS, rinsed in PBS and mounted in mounting medium with DAPI (Vector Laboratories, H-1200). Primary antibodies included mouse monoclonal anti-γ-tubulin (1:1000, Abcam ab11316), rabbit polyclonal anti-pericentrin (1:1000, Abcam ab4448) and rabbit monoclonal anti-pT73-Rab10 (1:1000, Abcam ab241060). Secondary antibodies were all from Invitrogen, were employed at a 1:1000 dilution, and included Alexa488-conjugated goat anti-mouse (Invitrogen, A11001) or goat anti-rabbit (Invitrogen, A11008), Alexa568-conjugated goat anti-mouse (Invitrogen, A11004) or goat anti-rabbit (Invitrogen, A11011) and Alexa647-conjugated goat anti-rabbit (Invitrogen, A21244).

For bromodeoxyuridine (BrdU) labeling of HEK293T, A549 and LCLs, cells were labeled with 10 μM BrdU (Abcam, ab142567) in respective full medium for 24 h, followed by fixation using 2%PFA/PBS for 20 min at room temperature. For PBMCs, cells were cultured in full medium for 48 h in either the presence or absence of PHA-M (1% v/v) with 10 μM BrdU for the last 24 h, followed by fixation as described above. In all cases, cells were washed with PBS for 10 min, permeabilized with 0.2% Triton-X100/PBS for 10 min at room temperature followed by DNA hydrolysis with 1 M HCl in PBS/0.2% Triton-X100 for 1 h at room temperature. Coverslips were rinsed three times with PBS and blocked for 1 h in 5% BSA (w/v) (Millipore, 126579) in 0.2% Triton-X100/PBS (blocking buffer). BrdU incorporation was detected by immunocytochemistry using an FITC-labelled BrdU antibody staining kit (BD, 556028) according to manufactureŕs conditions.

### Image acquisition and analysis

Images were either acquired on a Leica TCS-SP5 confocal microscope using a 63x 1.4 NA oil UV objective (HCX PLAPO CS) or on an Olympus FV1000 Fluoview confocal microscope using a 60x 1.2 NA water objective (UPlanSApo). Images were collected using single excitation for each wavelength separately and dependent on secondary antibodies, and the same laser intensity settings and exposure times were used for image acquisitions of individual experiments to be quantified. Around 13-16 image sections of selected areas were acquired with a step size of 0.5 μm, and maximum intensity projections of z-stack images analyzed and processed using Leica Applied Systems (LAS AF6000) image acquisition software or ImageJ. Only cells which displayed clear centrosomal staining were analyzed and in all cases, mitotic cells as determined by DAPI staining were excluded from the analysis. For both LCLs and PBMCs, 150-200 cells were quantified per sample, with nuclear diameter additionally determined for PBMC preparations. Sample processing and quantifications were performed blind to conditions. Some experimental conditions were independently quantified by an additional two observers blind to condition and at distinct research sites (Rutgers and Lille), with identical results obtained in all cases. Upon completion of all experiments, the patient code was unveiled for subsequent data analysis.

Quantification of the percentage of cells displaying pT73-Rab10 staining was performed over non-processed and non-saturated images acquired during the same time with the same laser intensities.

### Cell extracts and Western blotting

LCL cell clumps were dispersed by pipetting, and 1 x 10^6^ cells were treated in 1.5 ml tubes with 1 mM LLOMe (Sigma, L7393) or with DMSO or 50 nM MLi2 for 2 h at 37 °C.

Cells were centrifuged at 120 x g for 5 min at room temperature, resuspended gently in 1 ml PBS and then pelleted again. The cell pellet was resuspended in 100 μl PBS containing protease/phosphatase inhibitors and 1x SDS sample buffer. Samples were briefly sonicated three times, centrifuged for 10 min at 4 °C, and the supernatant boiled at 95 °C for 5 min. Alternatively, the cell pellet was resuspended in 100 μl freshly-prepared lysis buffer (50 mM Tris-HCl pH 7.5, 1% (v/v) Triton X-100, 1 mM EGTA, 1 mM Na_3_VO_4_, 50 mM NaF, 10 mM beta-glycerophosphate, 5 mM sodium pyrophosphate, 0.27 M sucrose, 0.1% (v/v) beta-mercaptoethanol, 1x cOmplete (EDTA-free) protease inhibitor cocktail (Roche, 04-693-124-001), 1 μg/ml Microcystin-LR (Enzo Life Sciences, Cat# number ALX-350-012-M001) and snap-frozen in liquid N_2_ and stored at – 80 °C. Protein concentration was estimated using the BCA assay (Pierce) according to manufactureŕs specifications. Extracts were mixed with SDS sample buffer supplemented with beta-mercaptoethanol (final volume 2.5% v/v) and heated at 95 °C for 5 min. Ten to fifteen micrograms of samples were loaded onto 4-20% precast polyacrylamide gels (Bio-Rad, 456-1096) and electrophoresed at 60 V (stacking) and 80 V (separating) in SDS running buffer (Tris-Glycine Running Buffer; 25 mM TRIS pH 8.6, 190 mM glycine, 0.1% SDS). Proteins were transferred to nitrocellulose membranes using the semi-dry Trans-Blot Turbo Transfer System (Biorad) for 10 min at constant 20 V (2.5 limit A). Membranes were blocked in TBS (20 mM Tris-HCl, pH 7.6, 150 mM NaCl) containing 50% of blocking buffer (Li-COR Biosciences, Intercept blocking buffer (TBS), 927-60001) for 1 h at room temperature, followed by cropping into three pieces for Li-COR multiplexing (top piece until 75 kD, middle piece until 37 kD, bottom piece). Membranes were incubated with primary antibodies in 50% TBST (TBS containing 0.1% (v/v) Tween-20) in 50% of Li-COR blocking buffer overnight at 4 °C. The top piece was incubated with a rabbit anti-S935-LRRK2 antibody (1:500, Abcam, ab133450) multiplexed with a mouse monoclonal anti-LRRK2 antibody (1:1000, Antibodies Inc, 75-235). The middle piece was incubated with a mouse monoclonal anti-α-tubulin antibody (1:10’000, Sigma, clone DM1A), and the bottom piece was incubated with a rabbit monoclonal anti-pT73-Rab10 antibody (1:1000, Abcam, ab230261) multiplexed with a mouse monoclonal total Rab10 antibody (1:1000, Sigma, SAB5300028). Membranes were washed three times for 10 min in 0.1% Tween-20/PBS, followed by incubation with secondary antibodies for 1 h at room temperature in 50% TBST in 50% LiCOR blocking buffer. Secondary antibodies included goat anti-rabbit IRDye 800CW and goat anti-mouse IRDye 680RD (1:10’000). Membranes were washed with 0.1% Tween-20/PBS for three times 10 min each. Blots were imaged via near-infrared fluorescent detection using Odyssey CLx imaging system, and quantification was performed using the instrument’s Image Studio software. For each LCL line, fresh extracts from 2-3 independent cultures were analyzed, and representative immunoblots are shown in the figures, with all immunoblots depicted in supplementary figures. All sample processing and quantifications were performed blind to conditions. Upon completion of all experiments, the patient code was unveiled for subsequent data analysis.

### Sequencing and data analysis

Whole exome sequencing (WES, Macrogen Korea) and long-range PCR and Sanger sequencing of the GBA gene were performed as previously described (*58, 84*). To identify potential genes/genetic variants that may mediate the MLi2-sensitive cohesion response, we examined the burden of rare (popmax AF < 0.0001 from gnomAD non-neuro population) mutations within subjects WES data. Both indels and SNVs were included in the analysis (multi-allelics were separated and indels were left-aligned). Variants previously identified within the ReFiNE full blacklist (0.01) were filtered. Only non-synonymous variants with functional refGene annotation in “splicing”, “exonic” or “exonic/splicing” were included. Gene burden analysis was performed using TRAPD (*85*) on all variants passing filters between TSS and TES. The UniProt (Apr 2020 hg38 release) of protein domain annotations were used for protein domain burden analysis. Rare variants were found within a total of 18,394 unique protein domains annotated to 4232 genes of which each were tested. For each gene/protein domain, a dominant and recessive test was performed. A dominant test uses cases with one or more variants within the gene of interest, whereas a recessive test uses 2 or more variants. The results from the burden test are summarized within **Table S1 and Table S2**. Note that none of the p-values pass false discovery rate correction (p<0.05, Benjamini & Hochberg). Gene ontology was performed on genes with a p-value < 0.05 (uncorrected) from each model using PANTHER. For pathway analysis, the autosomal dominant model/protein domains gene list was scanned for over-representation of KEGG pathways (http://www.webgestalt.org/) and also using the String Site (https://string-db.org/cgi/input?sessionId=bYAdyBStPoNM&input_page_show_search=on).

### Statistical analysis

Data were checked for normal distribution using the Shapiro-Wilk test. One-way ANOVA with Tukeýs post-hoc test was employed, with significance set at p < 0.05. All p values are indicated in the legends to figures. Spearman correlations were used to determine associations between protein levels and/or splitting values. Paired t-test analysis was performed for comparison of the splitting phenotypes in the presence versus absence of MLi2. All statistical analyses and graphs employed the use of Prism software version 9.5 (GraphPad, San Diego, CA).

## Data availability

Raw Western blot data and all sequencing data are available as supplemental figures and tables. All raw images of cohesion determinations are available upon request.

## Acknowledgements

We thank all patients and their families for donating blood samples, which made this study possible. We thank Laura Montosa, Besma Brahmia and Rollanda Bernadin for help with image aquisition, and Laurine Vandevinkel and Claire Deldycke for their help with cell culture. This work was funded by grants from the Michael J. Fox Foundation for Parkinso’s Research (MJFF-019358: SH, MCCH, JRM); MJFF-020338: ND). The PBMC Collection at Columbia University was funded by the Michael J. Fox Foundation (RNA).

## Author contributions

Conceptualization: SH, MCCH, JRM

Methodology: SH, MCCH, JRM

Investigation: YN, BF, RF, EF, IC, CL, JBK, EM, AD, JMT, RNA, ND, GH, YB, AV, CC

Visualization: SH, JBK, ND

Funding acquisition: SH, MCCH, JRM, ND, RA

Project administration: SH

Supervision: SH, MCCH, JRM, ND

Writing – original draft: SH

Writing – review and editing: YN, BF, RF, EF, IC, CL, JBK, EM, AD, JMT, SP, MD, RA, ND, GH, JRM, MCCH, SH

## Supplementary Material

**Figure S1.**
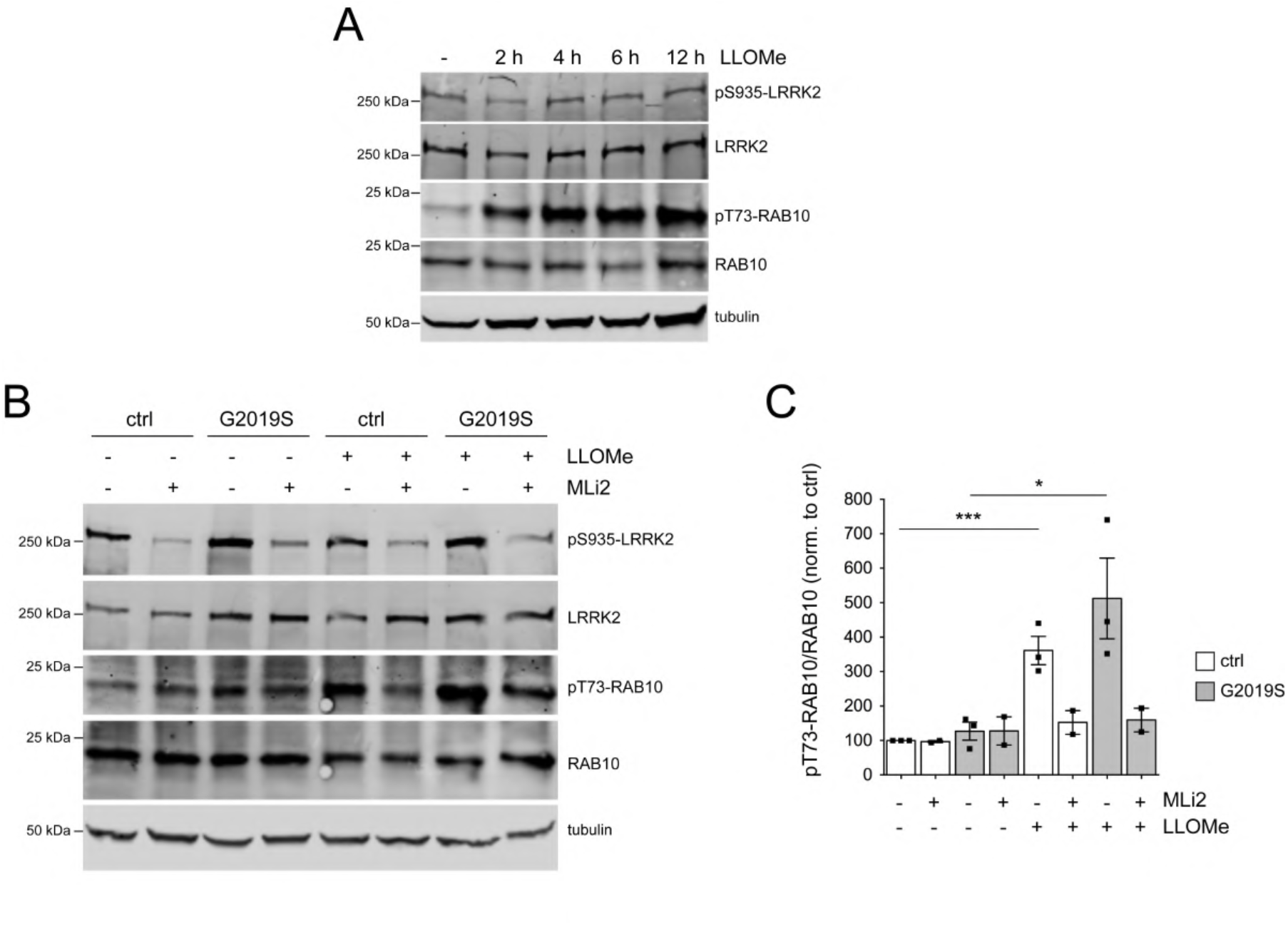
LLOMe treatment increases LRRK2-mediated pT73-Rab10 levels. (**A**) A control LCL line was treated with LLOMe (1 mM) for increasing amounts of time as indicated, and cell extracts were subjected to multiplexed immunoblot analysis. (**B**) A control and a *G2019S-LRRK2* LCL line were treated with LLOMe (1 mM) in either the absence or presence of MLi2 (50 nM) for 2 h as indicated before immunoblot analysis. (**C**) Quantification of pT73-Rab10/Rab10 levels from the type of experiments indicated in (B). N=3 independent experiments; ctrl versus ctrl + LLOMe (p = 0.003), G2019S mutation versus G2019S mutation + LLOMe (p = 0.032). ***p < 0.005; *p < 0.05.

**Figure S2.**
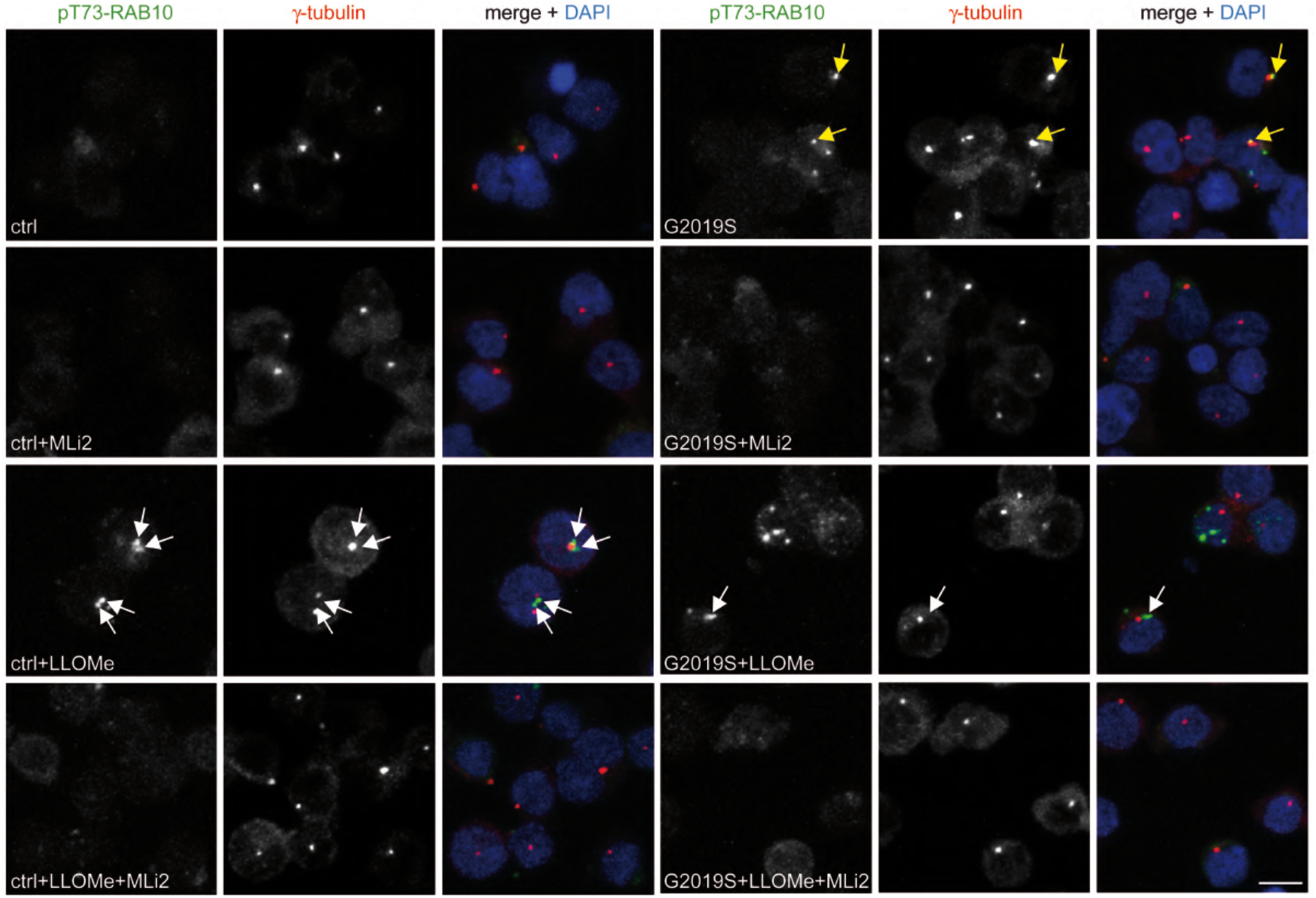
LLOMe treatment causes accumulation of pT73-Rab10 near centrosomes. Examples of healthy control (ctrl) or *G2019S-LRRK2* PD LCLs (G2019S) with or without treatment with LLOMe (1 mM) and MLi2 (50 nM) for 2 h as indicated, followed by staining with pT73-Rab10 antibody, γ-tubulin antibody and DAPI. Control cells display virtually no pT73-Rab10 staining. LLOMe treatment reveals dot-like pRab10 staining in some cells which is localized near the centrosome or in-between duplicated split centrosomes (white arrows). *G2019S-LRRK2* LCLs display pT73-Rab10 staining which is overlapping with a centrosomal marker in some cells (yellow arrows). LLOMe treatment reveals dot-like pT73-Rab10 staining near the centrosome in some *G2019S-LRRK2* cells (white arrows) similar to that observed in LLOMe-treated control cells. MLi2 treatment abolishes pT73-Rab10 staining in all cases. Scale bar, 10 μm.

**Figure S3.**
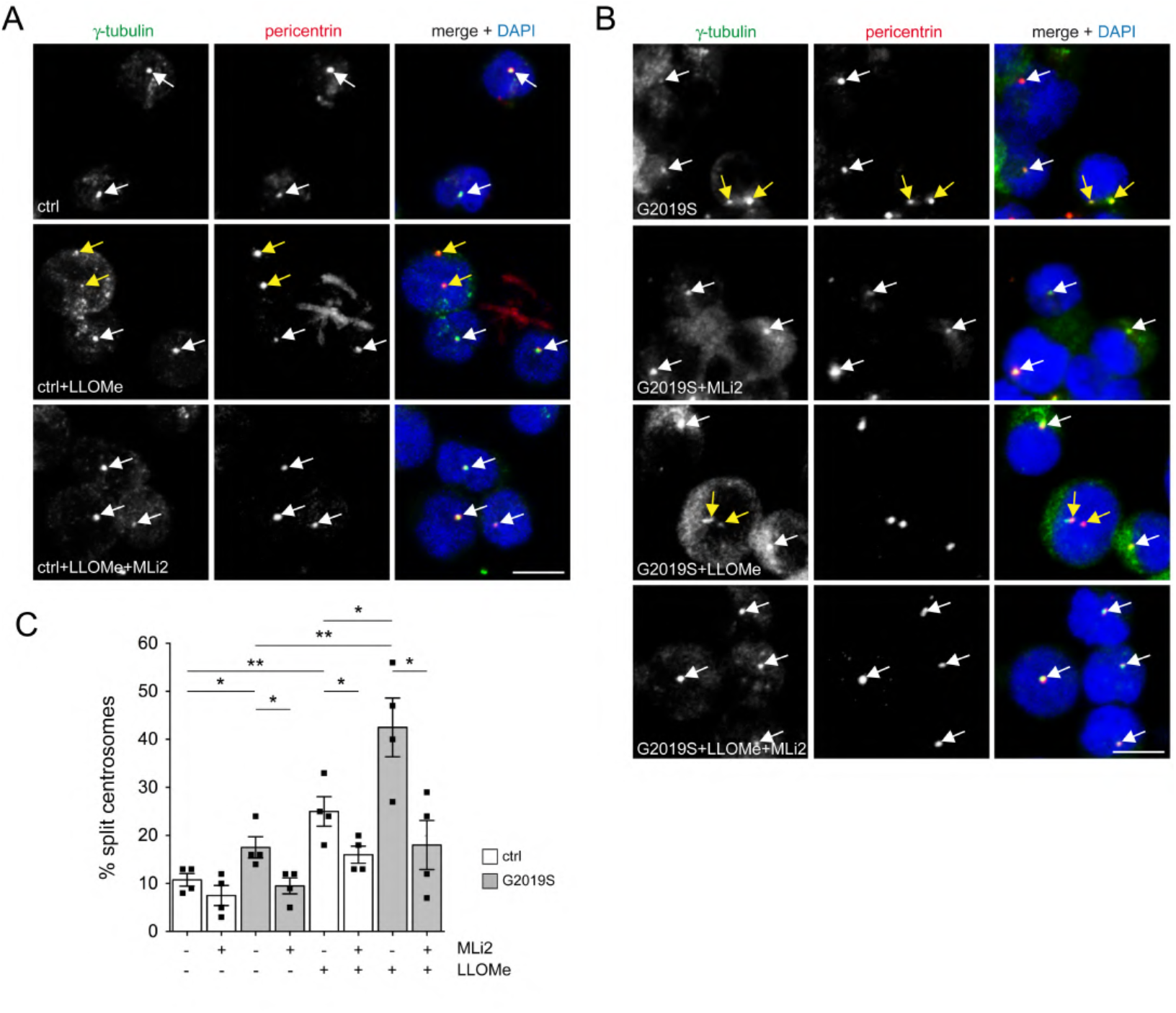
LLOMe treatment causes a LRRK2-mediated centrosomal cohesion deficit. (**A**) Examples of healthy control (ctrl) LCLs with or without treatment with LLOMe (1 mM) and MLi2 (50 nM) for 2 h as indicated, followed by staining with two centrosomal markers (γ-tubulin and pericentrin) and DAPI. Scale bar, 10 μm. White arrows point to centrosomes, yellow arrows point to duplicated split centrosomes. (**B**) Same as (A), but *G2019S-LRRK2* PD LCLs. (**C**) Quantification of the centrosome phenotype from control and *G2019S-LRRK2* PD LCLs treated as indicated. Bars represent mean ± s.e.m. (n=4 independent experiments); ctrl versus G2019S mutation (p = 0.039); ctrl versus ctrl + LLOMe (p = 0.005); G2019S mutation versus G2019S mutation + LLOMe (p = 0.008); ctrl + LLOMe versus G2019S mutation + LLOMe (p = 0.043); G2019S mutation versus G2019S mutation + MLi2 (p = 0.027); ctrl + LLOMe versus ctrl + LLOMe + MLi2 (p = 0.044); G2019S mutation + LLOMe versus G2019S mutation + LLOMe + MLi2 (p = 0.021); **p < 0.01; *p < 0.05.

**Figure S4.**
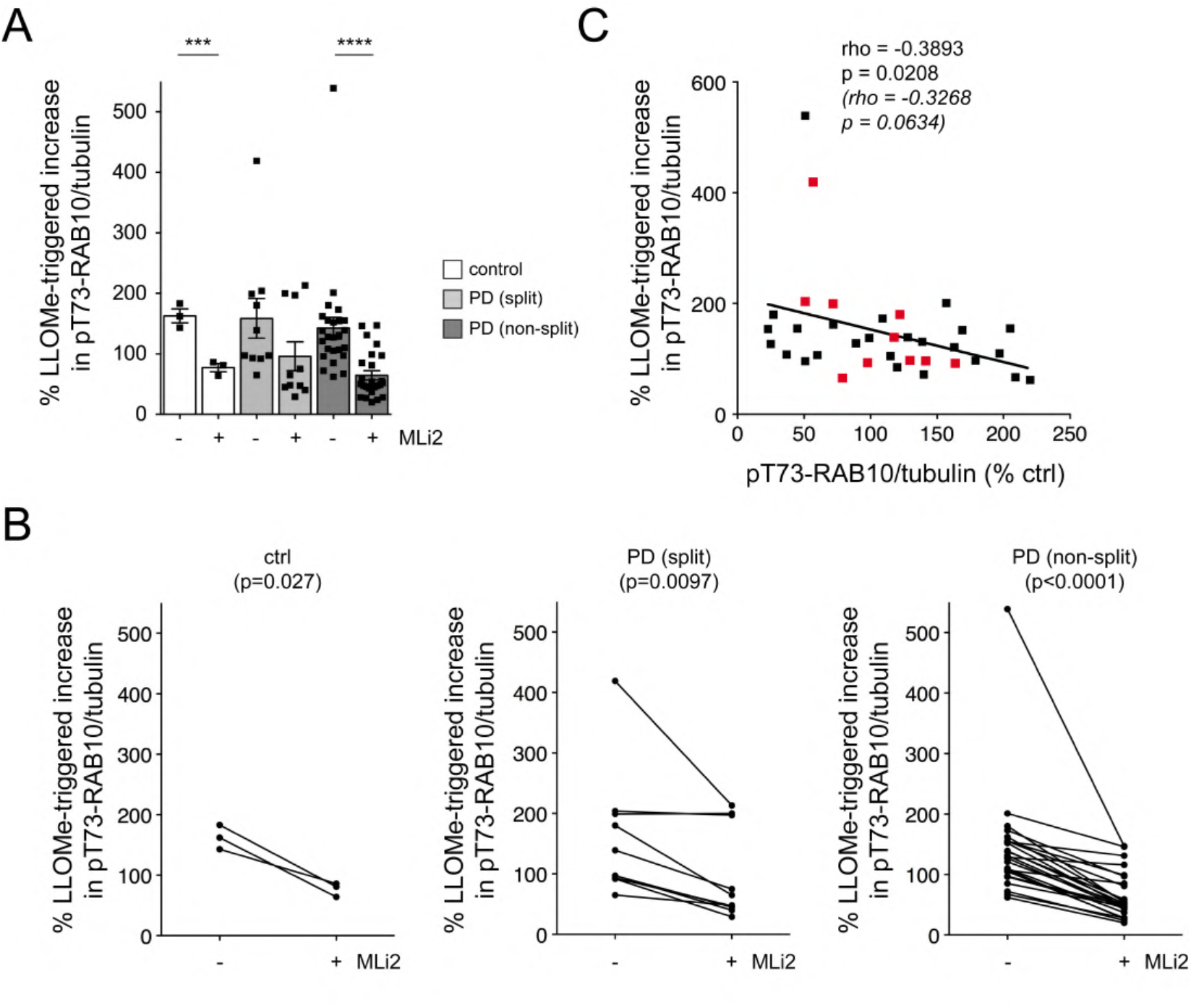
Effects of LLOMe treatment on LRRK2 kinase-mediated pT73-Rab10 levels. (**A**) The percentage of LLOMe-triggered increase in pT73-Rab10/tubulin levels in the absence or presence of MLi2 was calculated for each LCL line. Ctrl versus ctrl + MLi2 (p = 0.003); PD (non-split) versus PD (non-split) + MLi2 (p < 0.001). ****p < 0.001; ***p < 0.005. (**B**) Paired t-test analysis of LLOMe-triggered increase in pT73-Rab10/tubulin levels from each cell line in the absence or presence of MLi2. (**C**) Spearman correlation analysis between the percentage of LLOMe-triggered increase in pT73-Rab10/tubulin levels versus pT73-Rab10/tubulin levels in the absence of LLOMe treatment. There is a slight negative correlation between the basal pT73-Rab10/tubulin levels and the efficacy of the LLOMe-mediated increase in pT73-Rab10/tubulin levels. Red datapoints indicate the ten PD samples which display a centrosomal cohesion deficit. Rho and p values are indicated (in italics values without the two outliers).

**Figure S5.**
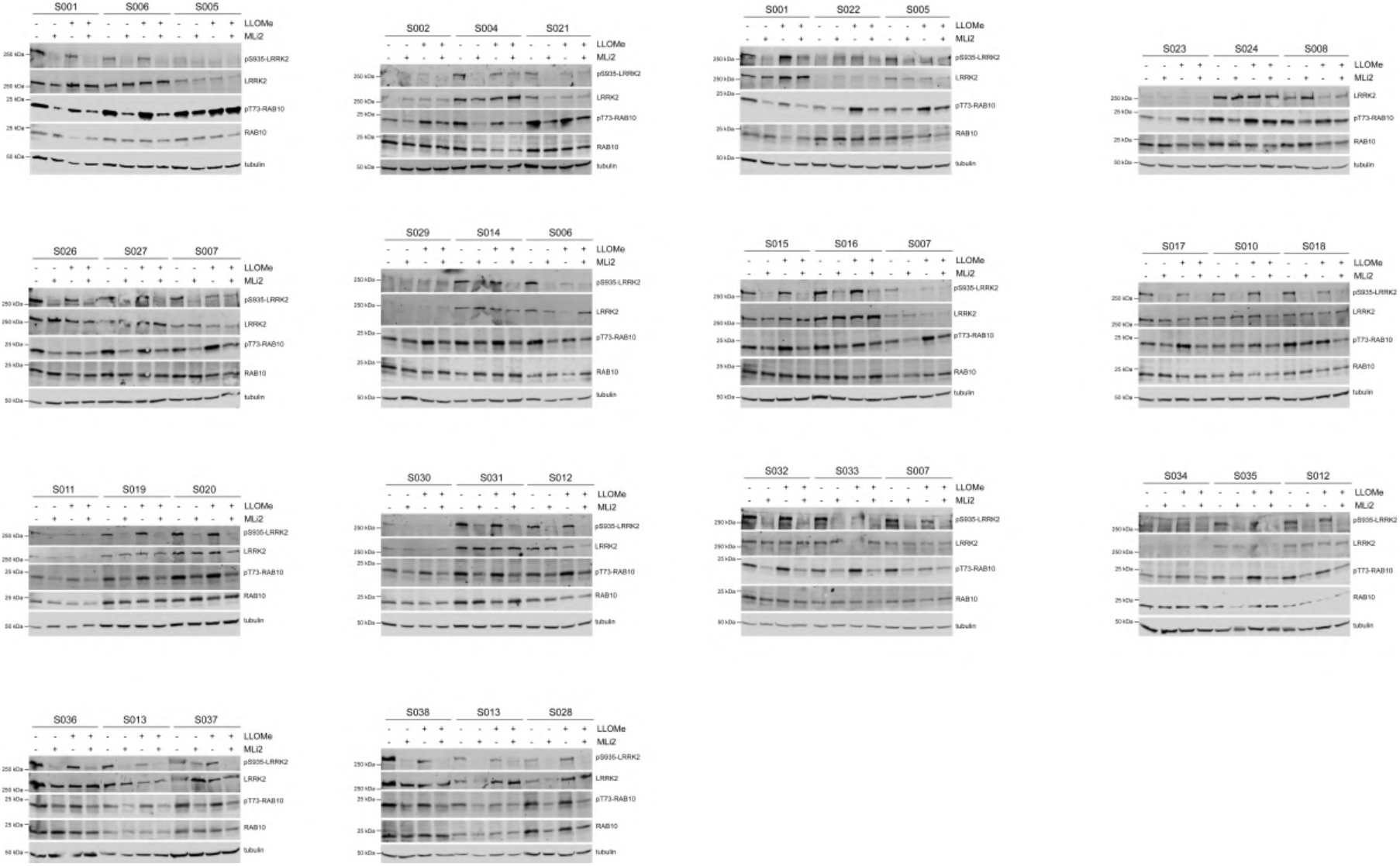
LCLs multiplexed quantitative immunoblot analysis for LLOMe-mediated and MLi2-sensitive changes in pT73-Rab10 phosphorylation. LCLs were treated with or without LLOMe (1 mM) and with either DMSO or MLi2 (50 nM) for 2 h as indicated prior to cell lysis. Fifteen μg of extracts were loaded and subjected to quantitative immunoblot analysis with the indicated antibodies, and membranes were developed using the Odyssey CLx scan Western blot Imaging system. pT73-Rab10 and total Rab10, as well as S935-LRRK2 and total LRRK2 antibodies were multiplexed. For each gel and cell line, the values of pT73-Rab10/Rab10 in the presence of LLOMe were then normalizad to those in the absence of LLOMe to obtain the % LLOMe-triggered increase in pRab10/Rab10 levels. Each line was processed in this manner 2-3 independent times.

**Figure S6.**
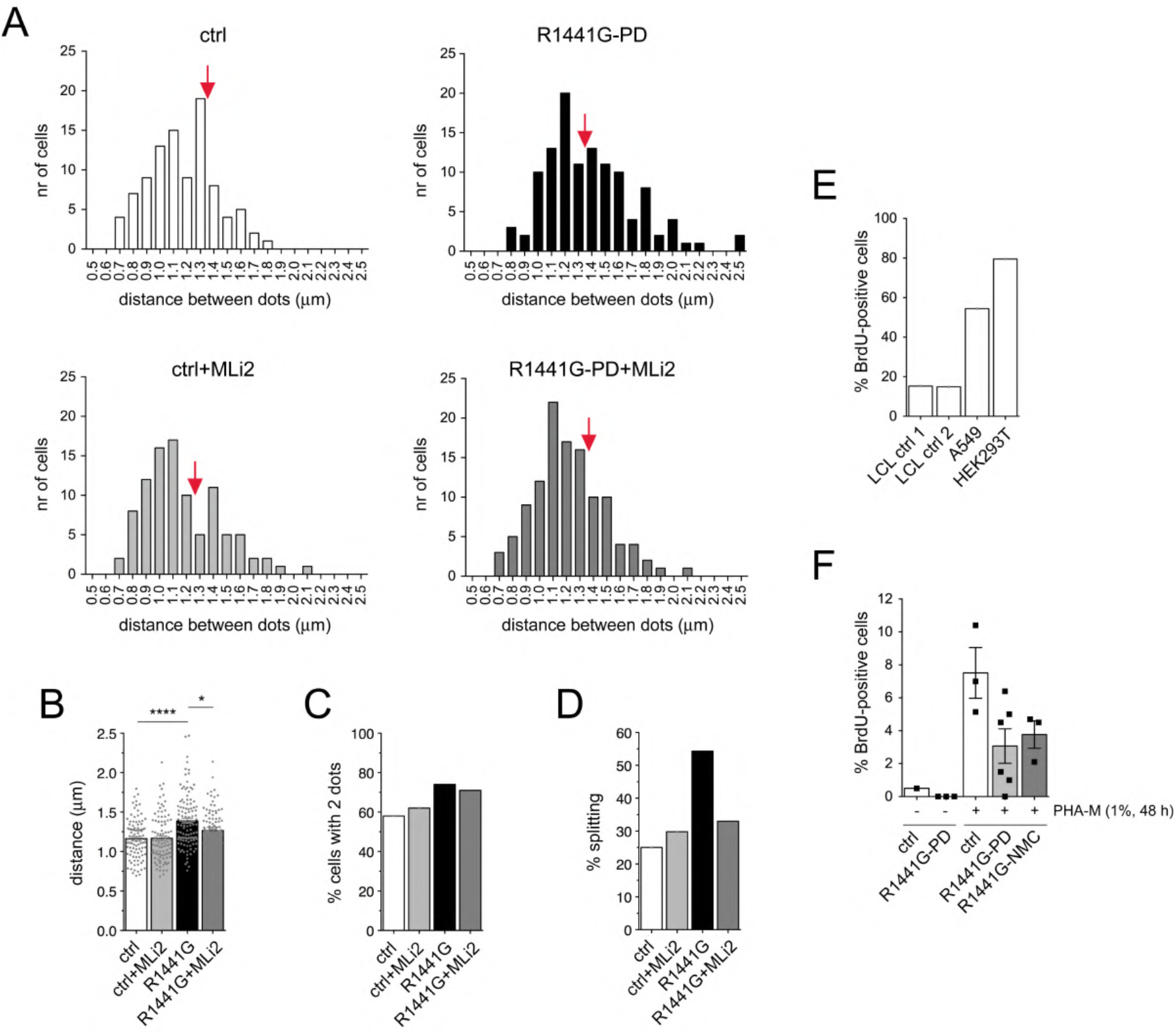
Cohesion analysis of pericentrin-positive dots from control and *R1441G-LRRK2* PD LCLs. (**A**) Frequency histogram distribution of the number of healthy control (left) and *R1441G-LRRK2* PD LCLs (right) displaying two pericentrin-positive dots in either the absence or presence of MLi2 (50 nM, 2 h) within binned distances as indicated. For each condition, around 100-120 cells displaying 2 pericentrin dots were quantified. (**B**) Mean distances between two pericentrin-positive dots in control and *R1441G-LRRK2* PD LCLs with or without MLi2 treatment. Ctrl versus R1441G mutation (p < 0.001); R1441G mutation versus R1441G mutation + MLi2 (p = 0.021). ****p < 0.001; *p < 0.05. (**C**) Percentage of cells displaying 2 pericentrin-positive dots in control and *R1441G-LRRK2* PD LCLs with or without MLi2 treatment. (**D**) Based on the frequency distribution shown in (A), cells were scored as having a C/C split phenotype when the distances between the two pericentrin-positive dots was > 1.3 μm (red arrow), and % splitting is indicated for control and *R1441G-LRRK2* PD LCLs with or without MLi2 treatment. (**E**) Two healthy control LCL lines, as well as wildtype A549 or HEK293T cells were treated with bromodeoxyuridine (BrdU)-containing media for 24 h before immunocytochemistry with an FITC-coupled anti-BrdU antibody and DAPI. BrdU-positive nuclei were scored from 200 cells each. (**F**) One healthy control (ctrl) and three *R1441G-LRRK2* PD PBMCs were labeled with BrdU-containing media for 24 h. Alternatively, three healthy controls, 6 *R1441G-LRRK2* PD and 3 *R1441G-LRRK2* NMC PBMCs were incubated in media containing the growth-enhancing mitogenic phytohemagglutinin-M (PHA-M, 1% v/v) for 48 h, with the last 24 h containing BrdU. Immunocytochemistry was performed as above, and BrdU-positive nuclei scored from 200 lymphocytes each.

**Figure S7.**
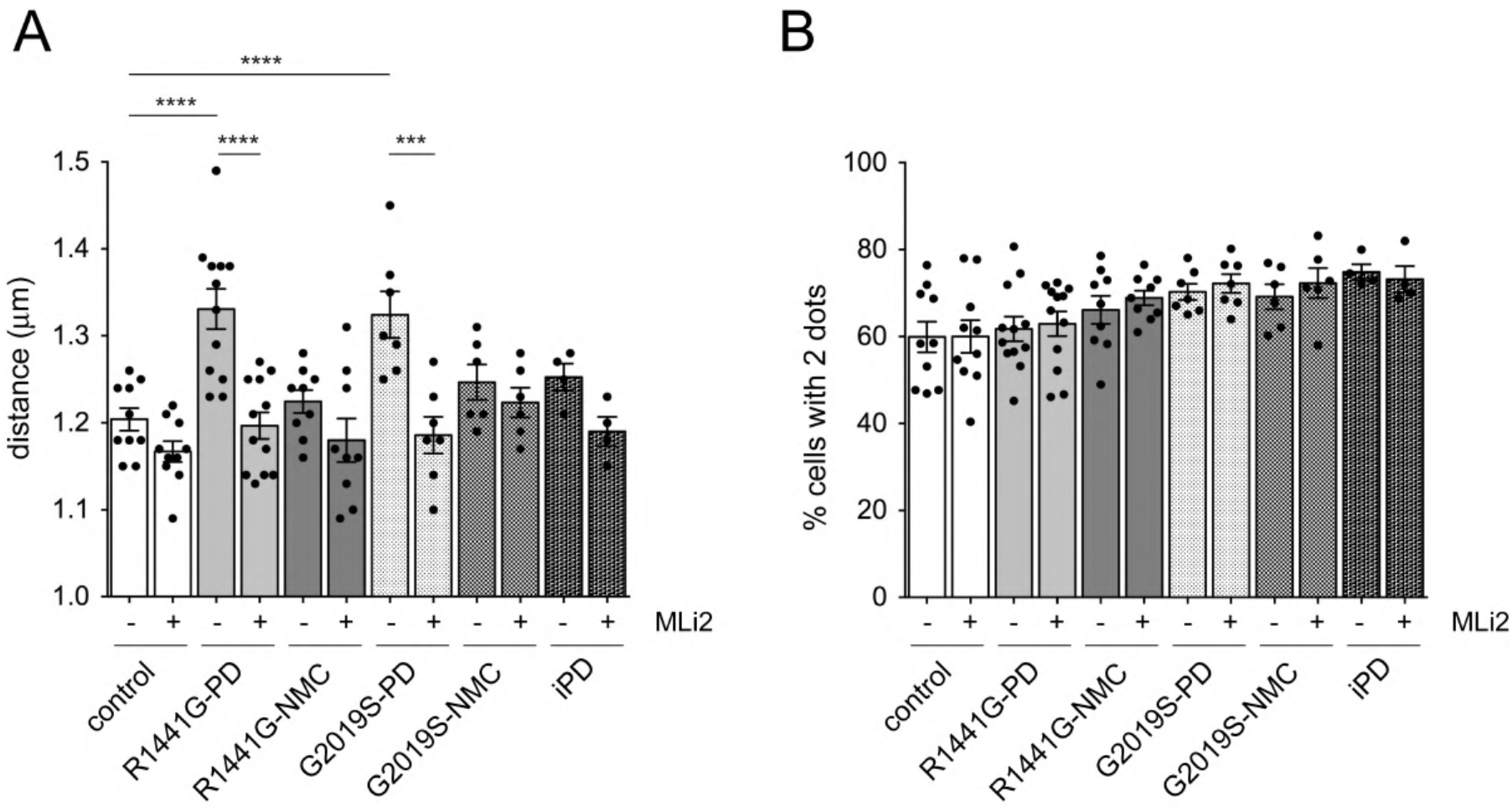
Percentage of cells with two pericentrin-positive dots and mean distances between dots in *R1441G-LRRK2* and *G2019S-LRRK2* LCLs. (A) Mean distance between the two pericentrin-positive dots from the different LCL lines. Ctrl versus R1441G mutation (p = 0.0002); ctrl versus G2019S mutation (p = 0.0004); R1441G mutation versus R1441G mutation + MLi2 (p < 0.0001); G2019S mutation versus G2019S mutation + MLi2 (p = 0.0015). ****p < 0.001; ***p < 0.005. (B) Quantification of the percentage of cells displaying two pericentrin-positive dots from a total of 100-150 cells per LCL line.

**Figure S8.**
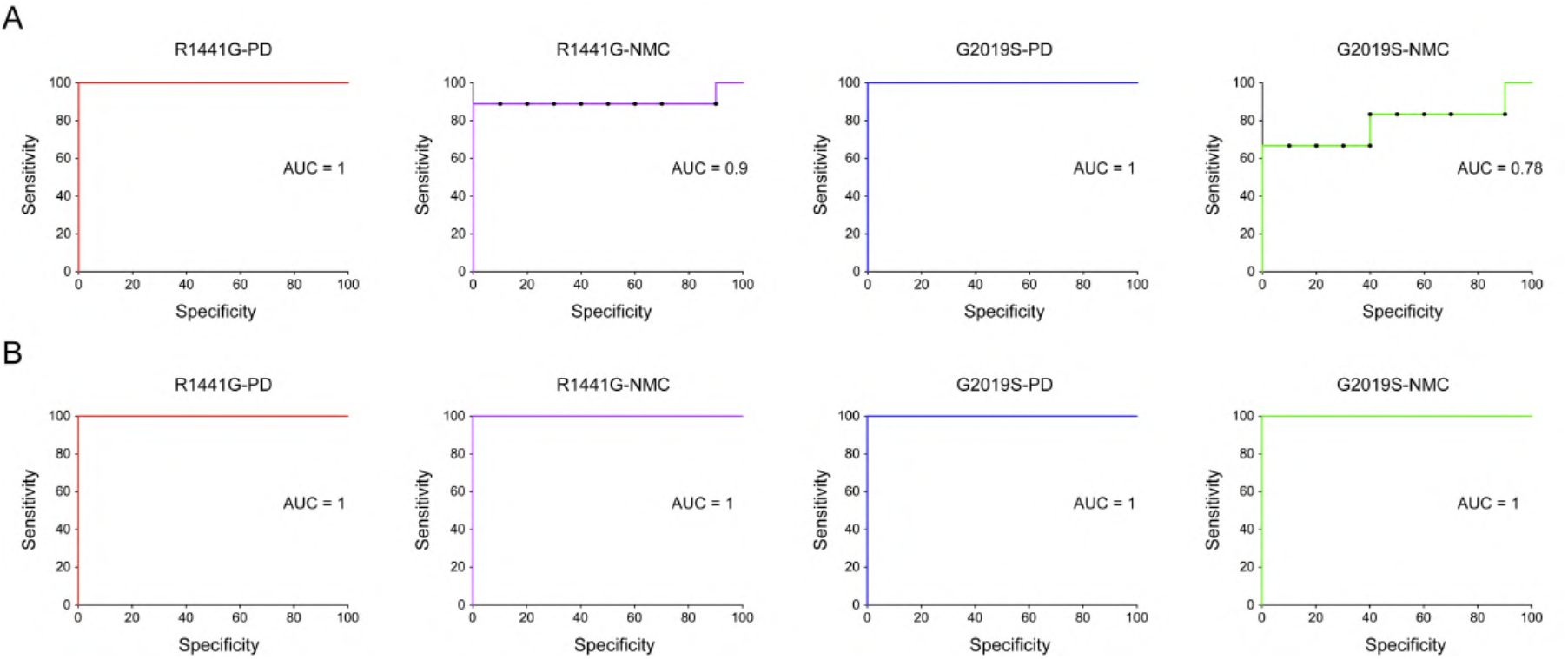
C/C cohesion phenotype and prediction AUC values for *LRRK2* mutation PD. (**A**) ROC curves showing prediction success for PD diagnosis in LCLs from *R1441G-LRRK2* PD, *R1441G-LRRK2* NMC, *G2019S-LRRK2* PD and *G2019S-LRRK2* NMC. (**B**) As in (A), but ROC curves for prediction success in PBMC lymphocytes.

**Figure S9.**
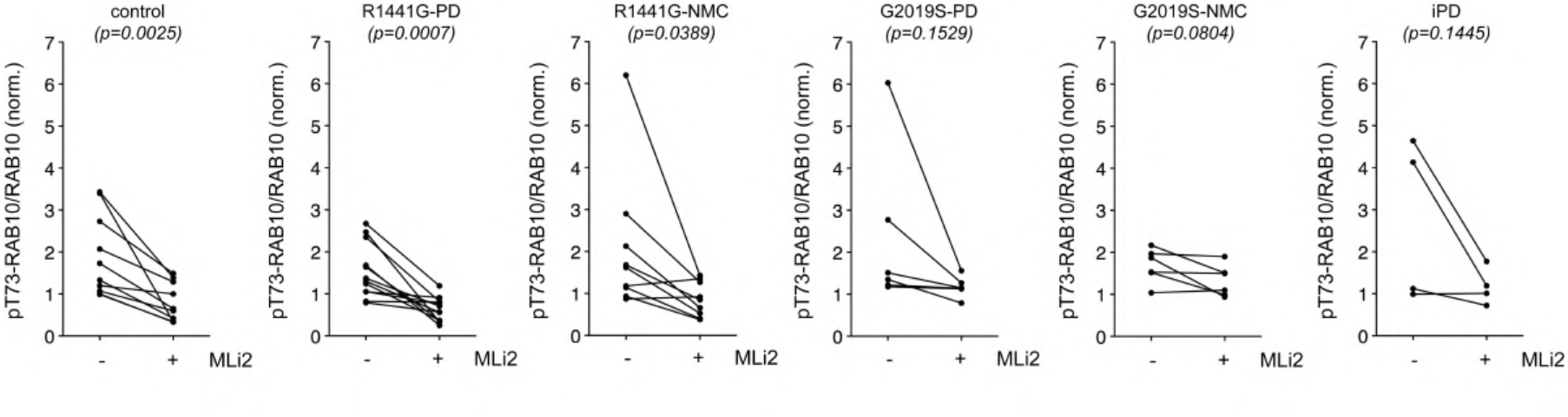
Analysis of MLi2-mediated alterations in pT73-Rab10/Rab10 levels in *R1441G-LRRK2* and *G2019S-LRRK2* LCLs. Paired t-test analysis of pT73-Rab10/Rab10 levels from each cell line in the absence or presence of MLi2. Note that the pT73-Rab10/Rab10 levels are reduced by MLi2 treatment in most cell lines.

**Figure S10.**
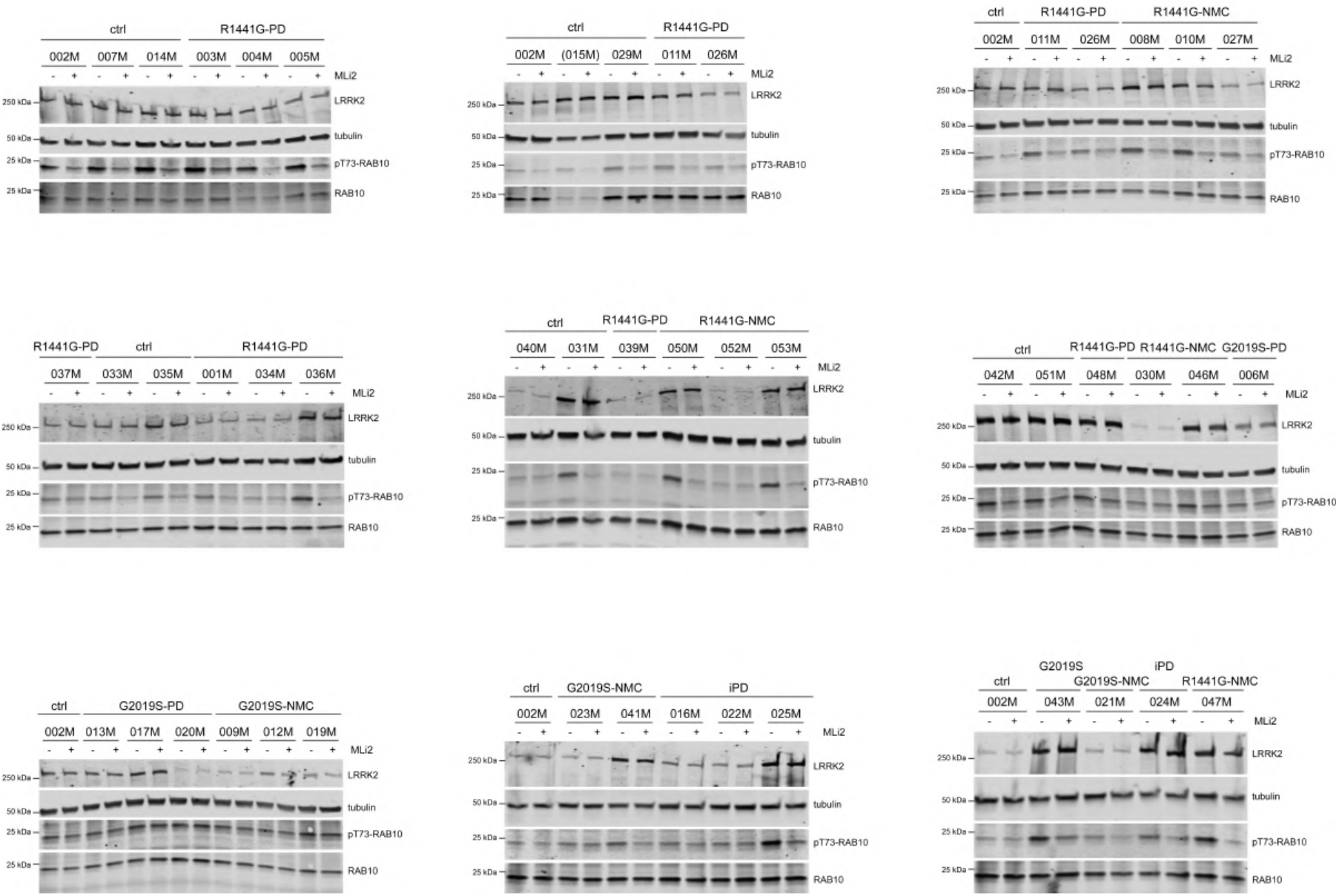
Multiplexed quantitative immunoblot analysis of LCLs. LCLs were treated with or without MLi2 (50 nM) for 2 h as indicated prior to cell lysis. Fifteen μg of extracts were loaded and subjected to quantitative immunoblot analysis with the indicated antibodies, and membranes were developed using the Odyssey CLx scan Western blot Imaging system. pT73-Rab10 and total Rab10 antibodies were multiplexed. For each gel and cell line, the values of pT73-Rab10/Rab10 were then normalized to an internal standard (control line 002M) to compare samples run on different gels. Each line was processed in this manner 2-3 independent times.

**Figure S11.**
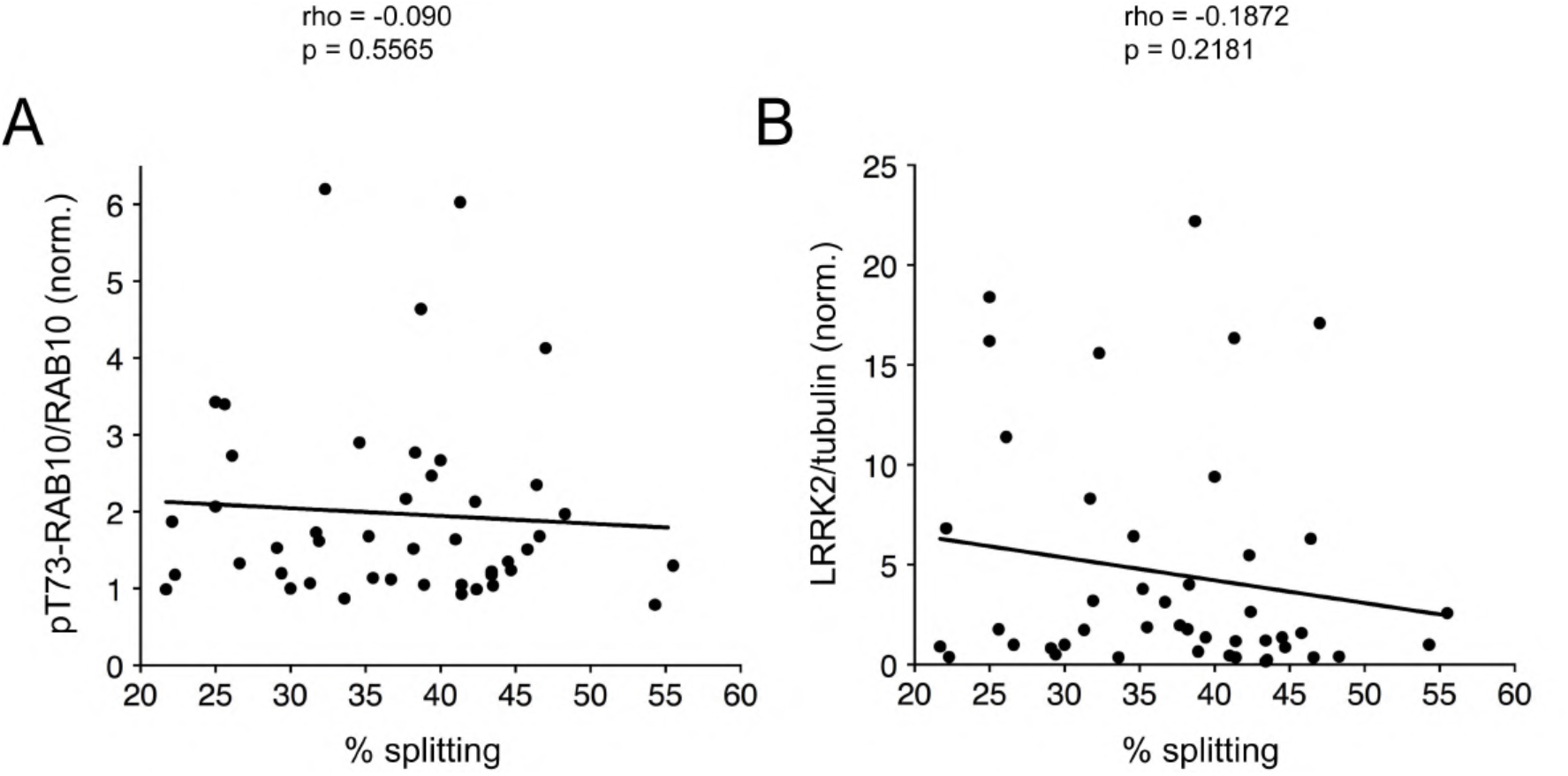
Correlation analysis between pT73-Rab10 or LRRK2 levels and the C/C cohesion phenotype. (**A**) Spearman correlation analysis between the levels of pT73-Rab10/Rab10 and the C/C cohesion phenotype (% splitting) from all LCL lines analyzed. (**B**) Spearman correlation analysis between the levels of LRRK2/tubulin and the C/C cohesion phenotype (% splitting) from all LCL lines analyzed. Rho and p values are indicated.

**Figure S12.**
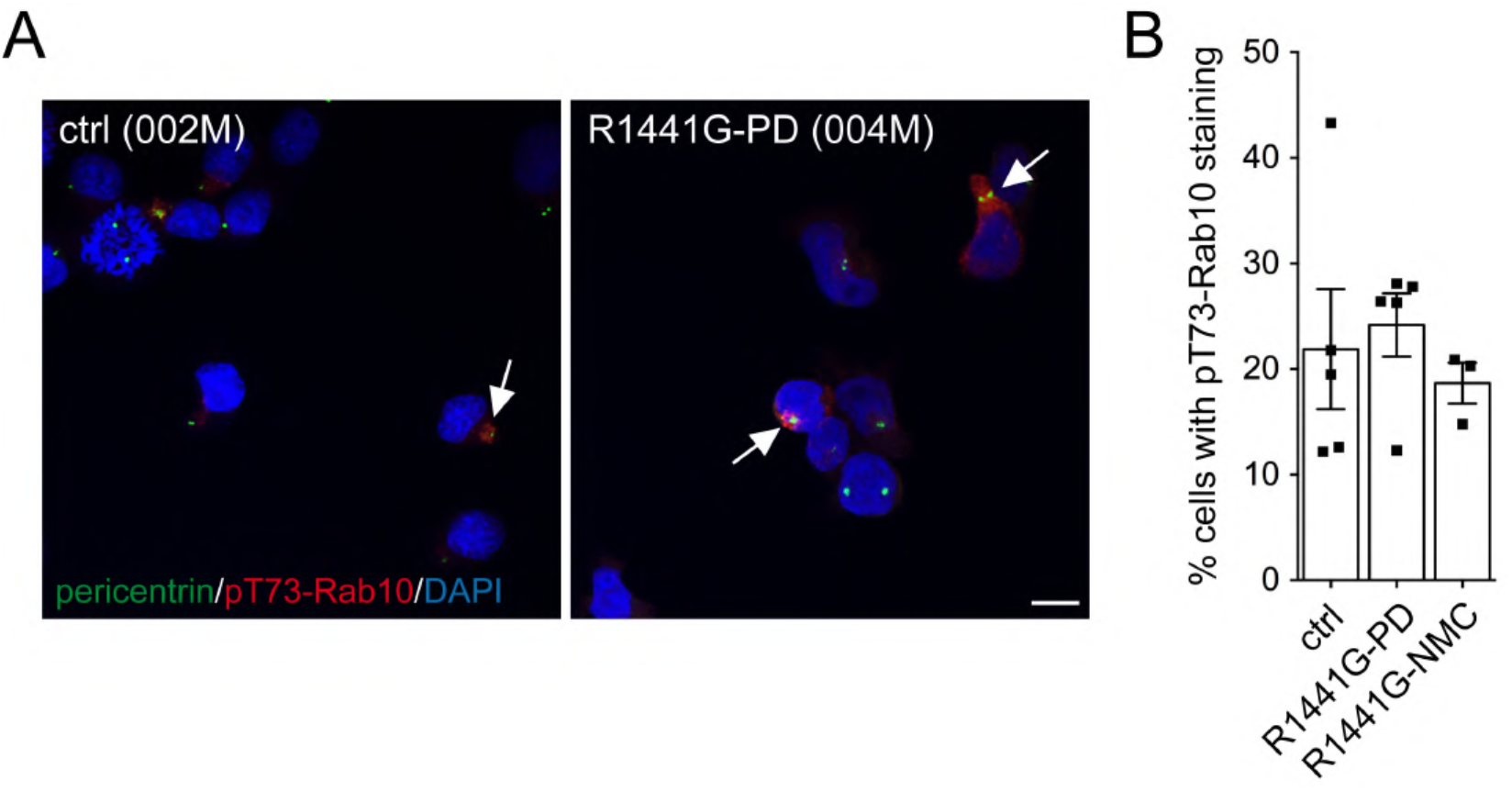
Quantification of the percentage of LCLs displaying pT73-Rab10 staining. (**A**) Representative images of control (ctrl) and *R1441G-LRRK2* PD LCLs stained with antibodies against pT73-Rab10 (red), pericentrin (green) and DAPI. Arrows point to cells displaying prominent pT73-Rab10 staining. Scale bar, 10 μm. (**B**) Quantification of the percentage of cells displaying pT73-Rab10 staining from 5 control, 5 *R1441G-LRRK2* PD and 3 *R1441G-LRRK2* NMC LCLs. Bars represent mean ± s.e.m.

**Figure S13.**
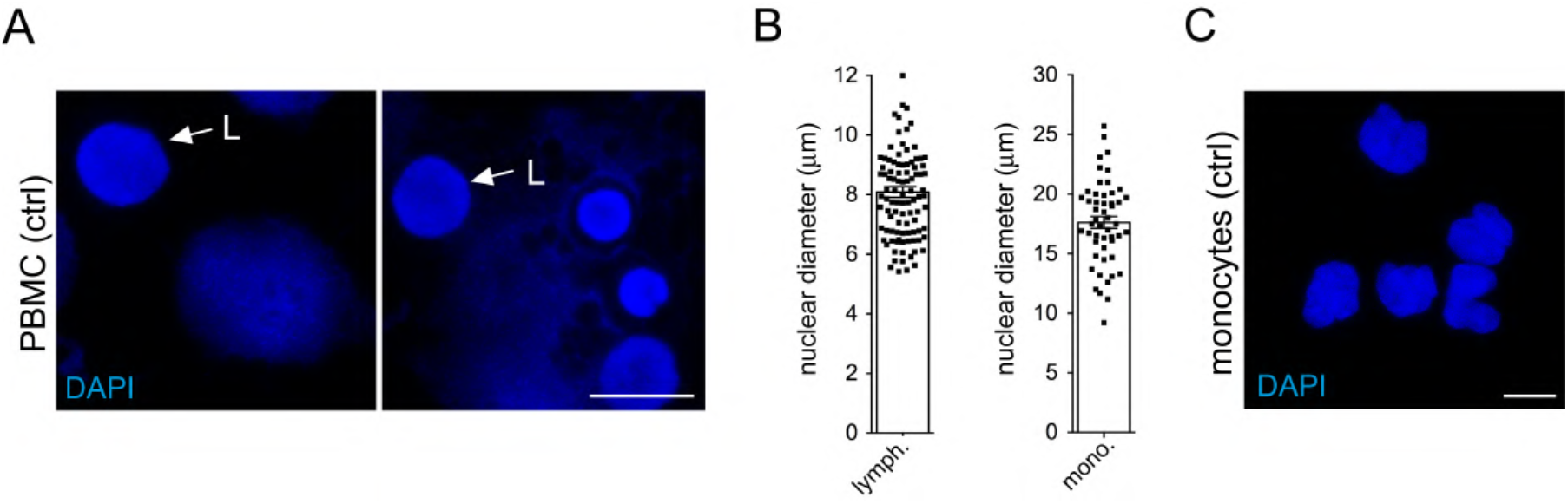
Detection of lymphocytes and monocytes in PBMC preparations. PBMCs were obtained from the LRRK2 Biobanking Initiative from five healthy controls and five *G2019S-LRRK2* PD patients. PBMCs from the same patients had been purified either employing sodium citrate-coated or heparin-coated tubes and had been cryopreserved at 3 x 10^6^ cells/tube. Cell debris was observed in both preparations, but was particularly extensive in PBMCs purified with sodium citrate-coated tubes. Therefore, analysis was only performed for PBMCs purified with heparin-coated tubes. Example of control PBMCs stained with DAPI. L, lymphocytes. Scale bar, 10 μm. Average nuclear diameter of lymphocytes (left) (n=98 cells) and monocytes (right) (n=51 cells) from healthy control PBMC preparation as shown in (A). (**C**) Purified monocytes were stained with DAPI. Note the larger and typically kidney-shaped nucleus. Scale bar, 10 μm.

**Figure S14.**
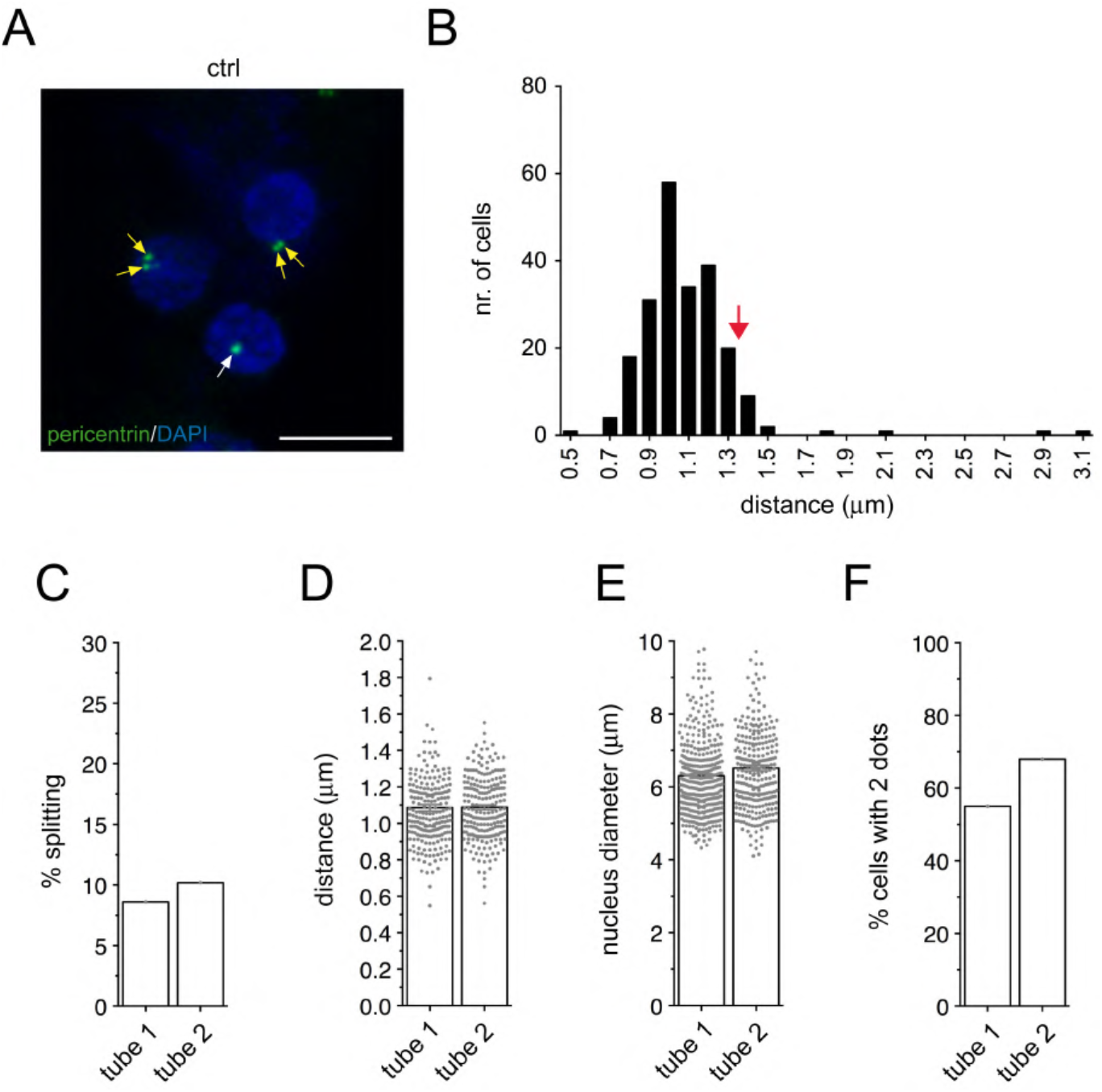
C/C cohesion determination and test-retest reliability in lymphocytes from control PBMC preparation. (**A**) Example of lymphocytes from healthy control (ctrl) PBMC preparation stained for pericentrin and DAPI. White arrow points to pericentrin-positive structure, yellow arrows point to two pericentrin-positive structures with or without a splitting phenotype. Scale bar, 10 μm. (**B**) Frequency histogram distribution of the number of healthy control lymphocytes displaying two pericentrin-positive dots, with binned distances as indicated. Around 200 cells displaying 2 pericentrin dots were quantified. (**C**) Mean distance between two pericentrin-positive dots. (**D**) Mean nucleus diameter for lymphocytes analyzed. (**E**) Percentage of cells displaying two pericentrin-positive dots. (**F**) Based on the frequency distribution shown in (B), cells were scored as having a split phenotype when the distances between the two pericentrin-positive dots was > 1.3 μm (red arrow), and % splitting is indicated. Two independent cryopreserved PBMC tubes were analyzed one month apart, with excellent test-retest reliability within an individual over time.

**Figure S15.**
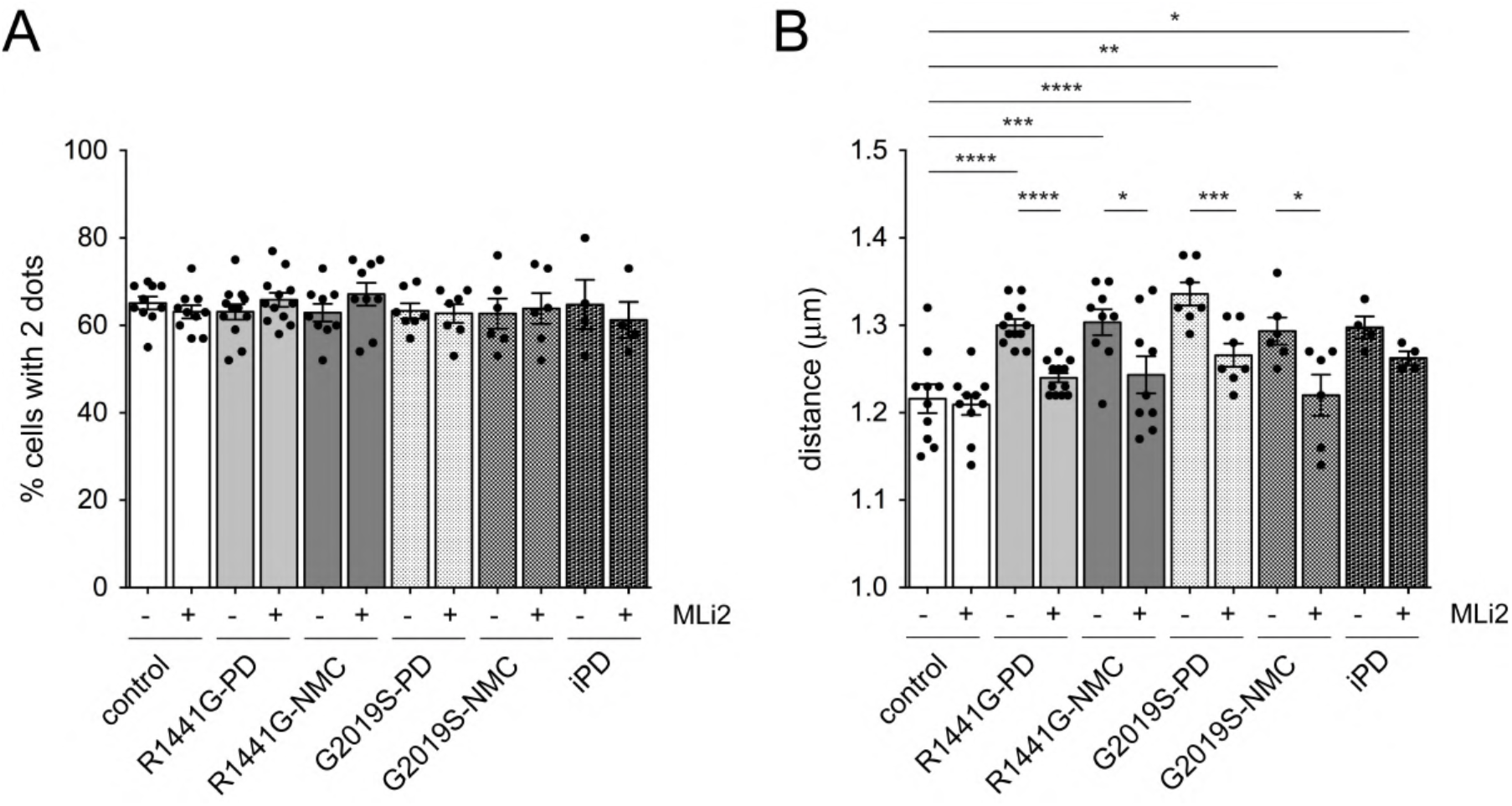
Percentage of cells with two pericentrin-positive dots and mean distances between dots in *R1441G-LRRK2* and *G2019S-LRRK2* lymphocytes. (A) Quantification of the percentage of cells displaying two pericentrin-positive dots from a total of 100-150 lymphocytes per PBMC preparation. (B) Mean distance between the two pericentrin-positive dots from lymphocytes of each PBMC preparation. Ctrl versus R1441G mutation (p < 0.0001); ctrl versus R1441G NMC (p = 0.001); ctrl versus G2019S mutation (p < 0.0001); ctrl versus G2019S NMC (p = 0.007); ctrl versus idiopathic PD (p = 0.0125); R1441G mutation versus R1441G mutation + MLi2 (p < 0.0001); R1441G NMC versus R1441G NMC + MLi2 (p = 0.034); G2019S mutation versus G2019S mutation + MLi2 (p = 0.0028); G2019S NMC versus G2019S NMC + MLi2 (p = 0.0266). ****p < 0.001; ***p < 0.005; **p < 0.01; *p < 0.05.

**Figure S16.**
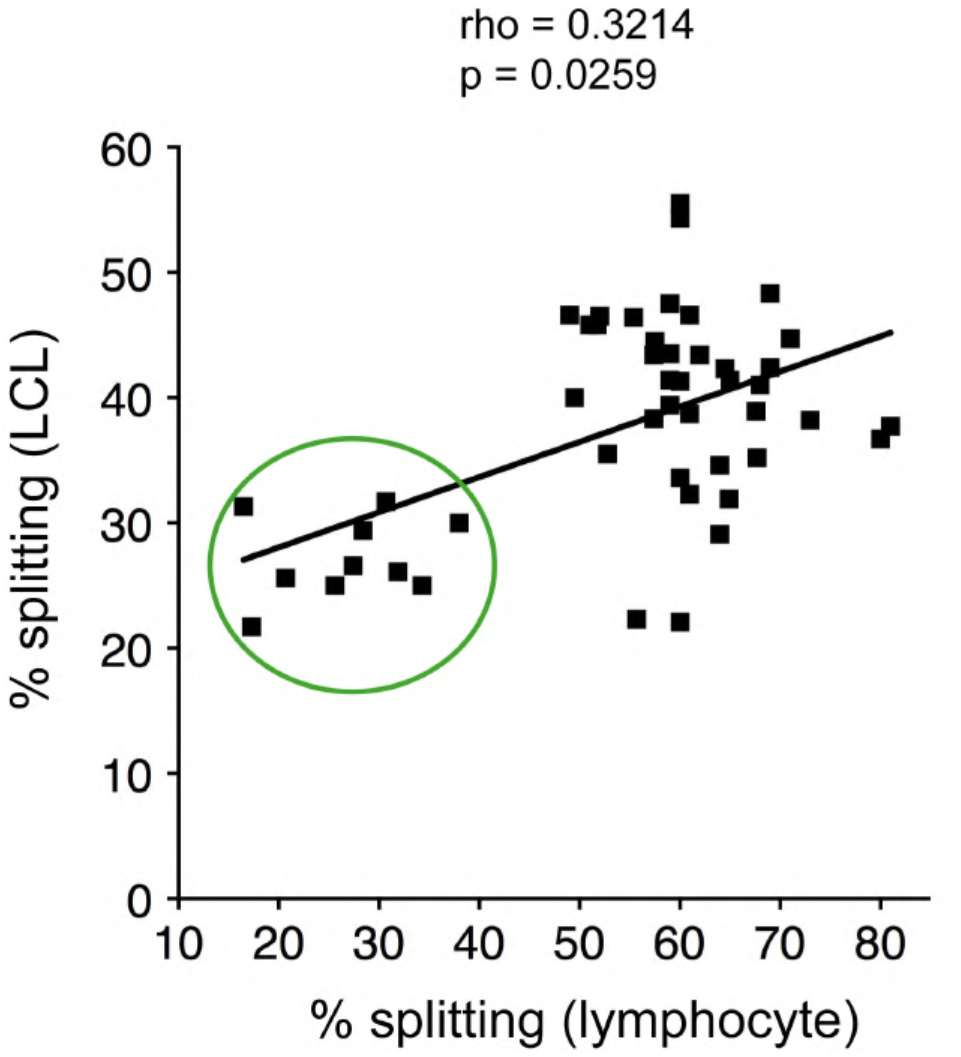
C/C cohesion phenotype in LCLs versus lymphocytes from healthy control and *LRRK2* mutation carriers. Spearman correlation analysis between the percentage of C/C splitting observed in lymphocytes and in immortalized LCLs from each patient. Healthy control patients are circled in green.

